# Modeling gene regulatory perturbations via deep learning from high-throughput reporter assays

**DOI:** 10.64898/2026.03.27.714770

**Authors:** Revathy Venukuttan, Richard Doty, Alexander Thomson, Yutian Chen, Boyao Li, Yuncheng Duan, Alejandro Barrera, Katherine Dura, Kuei-Yueh Ko, Hilmar Lapp, Timothy E. Reddy, Andrew S. Allen, William H. Majoros

## Abstract

Assessing likely variant effects on phenotypes is of critical importance in diagnostic settings, and while much progress has been made in interpreting genic mutations based on our understanding of coding sequence, noncoding variants can be much more challenging to reliably interpret based on DNA sequence alone. High-throughput reporter assays such as STARR-seq and MPRA have shown utility in experimentally measuring regulatory effects of noncoding variants present in samples but provide no readout for variants not present in the assay inputs. However, whole-genome reporter assays provide copious data that can be used to train predictive models for prioritizing variants not directly observed in the experiment. We describe a retrainable predictive modeling framework, BlueSTARR, for this task, and present results of training several models with this framework on whole-genome STARR-seq data from two cell lines and one drug treatment. Using these models, we uncover a global signature across the human genome consistent with purifying selection against both loss-of-function and gain-of-function regulatory variants, with the latter showing a significant bias consistent with selection against gains of cis regulatory function in closed chromatin proximal to genes. By testing the model on synthetic enhancers with binding motifs for transcription factors GR and AP-1, we find that when trained on drug perturbation data, the model is able to learn distance-dependent and treatment-dependent binding patterns and their resulting reporter gene activation. These results demonstrate that lightweight, easily retrainable models such as ours have utility in probing latent signals present in novel experimental data. Finally, we find only modest differences in performance between different deep-learning architectures when trained on this single data modality, and while somewhat greater predictive accuracy can be achieved with much larger models trained at great expense on many terabytes of data, there is still copious room for improvement even for industrial strength, state-of-the-art models.

## 1. Introduction

While the majority of currently known human disease-causing mutations have been found within protein-coding genes [1], association studies routinely uncover signals strongly implicating the roughly 98 percent of the genome that is annotated as noncoding, and it has been speculated that the majority of actual causal variants in disease may in fact be noncoding [2], [3], [4]. Because the noncoding genome is still so poorly understood compared to protein-coding genes, interpreting the likely effects of noncoding variants remains very challenging, making genetic diagnosis based on noncoding variants much more difficult than for coding variants. Given that majority of disease-associated mutations appear to be noncoding, there is a critical and yet unmet need to improve our ability to interpret noncoding variants.

High-throughput reporter assays such as *Self Transcribed Active Regulatory Region* (STARR-seq) and *Massively Parallel Reporter Assay* (MPRA) have shown great utility in experimentally measuring regulatory effects of noncoding variants, predominantly via episomal reporter constructs. By cloning genomic regions out of the endogenous genome and into the episomal reporter, relative allelic effects can be measured outside their native genomic context, thereby bypassing confounding issues such as local chromatin state. However, such assays provide no direct readout for variants that were either not present in the input samples, or were lost during library preparation or sequencing, due to allelic dropout. While including multiple samples simultaneously in one assay increases the number of variants that can potentially be tested at once [5], substantially expanding the number of samples increases the risks of dropout for rare variants [6], which are often the variants of greatest interest in disease studies. Moreover, while these assays have to date been used to assay tens of millions of variants simultaneously, those efforts have resulted in experimental characterization of only a miniscule proportion of the total noncoding genomic space.

With the rise of modern deep-learning approaches for predictive modeling of the human genome [7], [8], [9], [10], it is natural to consider the potential for using data from high-throughput reporter assays for training predictive models of noncoding variant effects. Indeed, previous studies have demonstrated that whole-genome STARR-seq data from model organisms such as Drosophila melanogaster can be used to train neural network models for making detailed predictions of the impact of noncoding variants on transcriptional regulation [11]. As whole-genome STARR-seq experiments applied to human samples can generate billions of unique fragments [12], these experiments can provide potentially very rich data sets for training useful predictive models of human cis regulatory logic.

Here we describe efforts aimed at constructing and training such models using data produced from whole-genome STARR-seq experiments performed on human K562 (human erythroleukemic cells) and A549 (human adenocarcinomic alveolar basal epithelial cells) cell lines under either control (DMSO/vehicle) or synthetic glucocorticoid drug treatments. While more complex predictive models trained on much larger and substantially more diverse training data have recently been described [7], [8], [9], [10], here we demonstrate the utility of more lightweight single-modality models in probing gene regulatory biology and signals of evolutionary constraint.

In particular, we show that data from custom experiments involving specific perturbations such as drug treatments can be used to train models that are able to learn and approximately reconstruct nuanced patterns of regulatory syntax and drug response. We additionally use the trained models to explore the hypothesis that gain-of-function variants in noncoding regions not currently annotated as regulatory elements may have the potential to be disruptive and subject to natural selection. In light of the latter result, we advocate for the use of such predictive models of noncoding variant effects to systematically scan for potential gain-of-function variants outside of known regulatory elements, as these may represent an important class of currently under-characterized disease mutations.

## 2. Materials and Methods

### 2.1 Data pre-processing

Self-Transcribing Active Regulatory Region sequencing (STARR-seq) is a reporter assay that measures genome-wide enhancer activity [13]. Candidate regulatory sequences are cloned downstream of a minimal promoter such that active enhancers drive their own transcription. The abundance of RNA transcripts obtained from the assay reflects enhancer activity and sequencing of the plasmid DNA provides a measure of input abundance.

Whole-genome STARR-seq experiments were performed in K562 cells [12], as well as in A549 cells treated with either dexamethasone (“A549/DEX”), a potent synthetic glucocorticoid drug, or with DMSO vehicle (“A549/DMSO”), a control [14]. The K562 experiment included 3 replicates of input DNA and output RNA. The A549 experiments included 5 replicates of input and 4 replicates of output. Data processing was performed by a publicly available workflow pipeline implemented in Common Workflow Language (CWL) [15].

DNA and RNA read counts were processed to generate genomic windows along with associated sequence and count information. The whole genome sequence from the input library was divided into overlapping windows of 300 bp with a 50 bp step size. Read coverage within those windows was quantified using bedgraph [16] files and filtering was performed on the sum of read counts in these bins across replicates. Windows with total DNA read counts exceeding a threshold of 100 reads were retained. This step ensured sufficient input coverage and removed regions with low signal. To prevent potential data leakage caused by highly similar genomic sequences, we removed paralogous sequences using BLASTN. Sequences sharing greater than 90% sequence identity and at least 100 identical consecutive bases were identified and removed. This filtering step removed approximately 60% of sequences in the A549 dataset and 8% in the K562 dataset.

For each retained genomic window, the corresponding DNA sequence was extracted from the GRCh38 reference genome using bedtools [17], producing sequence-count pairs suitable for model training. Sequences were then converted into one-hot encoding where each nucleotide is represented as a four-digit binary number with one 1 and three 0 digits. RNA and DNA counts from the selected genomic windows were normalized for replicate-specific library sizes by dividing by the total number of reads in each library. Following normalization, enhancer activity for each window was estimated using a naive estimator defined as:

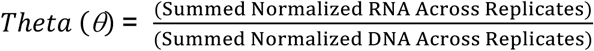

These activity estimates were used as target values for training the neural network. For experiments described here, models were by default trained on 1.6 million sequences, totaling 480 Mb of sequence data for the models taking 300 bp inputs.

### 2.2 Model architectures and implementation

BlueSTARR was implemented by extending the DeepSTARR model [11] and permits easy modification of the overall neural network architecture via a simple configuration file, allowing for rapid comparison of different model architectures. The default architecture consists of a convolutional neural network (CNN) that accepts one-hot encoded 300 bp DNA sequences as input. Our default model comprises five one-dimensional convolutional layers with 1024, 512, 256, 128 and 64 filters, and kernel sizes of 8,16, 32, 64 and 128, respectively (Suppl. Fig. S9). This choice was made to maintain a large receptive field without using pooling after each layer (as is done in DeepSTARR). Each convolutional layer is followed by batch normalization and ReLU activation, and a dropout rate of 0.5 is applied before each convolution. All layers use the same padding with no dilation, no residual connections and no intermediate pooling. The final convolutional layer outputs were aggregated by global average pooling and connected directly to a single output neuron for prediction. The model was trained using the Adam optimizer [18] with a learning rate of 0.002, a mean-squared error (MSE) loss function, batch size of 128, and an early stopping “patience” value of 10. All the aforementioned hyperparameters can be easily adjusted via edits to a simple configuration text file.

Variations of the above-mentioned architecture were explored to evaluate different receptive fields, sequence lengths, number of layers, inclusion of attention mechanism, and a custom loss function based on negative log-likelihood (NLL). Details of these modifications are described in the Supplementary Methods Section S1.2. To enable the easy incorporation of such changes to the base architecture, the code has been packaged as a flexible, easy to use software package called BlueSTARR. BlueSTARR is implemented in Python version 3.10+ using the Keras v2 and TensorFlow v2.15 libraries for building, training and evaluating the model. The software is available for download from GitHub as an open-source software under the MIT license.

### 2.3 Data preparation for model training

To construct the training, test and validation sets, we applied two downsampling strategies. In unbiased downsampling, each sequence was accepted into the sample with a fixed probability N/M, where N is a desired sample size and M is the total number of records. Whereas in biased downsampling, the naive theta distribution was divided into B equal width bins, and per-bin acceptance probabilities were computed as min (1, N/(M*B*p_i_)) where p_i_ is the observed proportion of records in bin *i*.

The initial preprocessing steps (described in section 2.1) generated 19 million training examples for the K562 STARR-seq data, 6 million training examples for the A549/DMSO STARR-seq data and 7 million training examples for A549/DEX STARR-seq data. Since training the model with full data can be computationally expensive, we conducted an experiment which trained the model on 0.8M, 1M, 1.3M, 1.5M and 1.55M training examples. These models were then benchmarked against the Kircher et al MPRA data [19] and AUROC values were visualized for each model (Fig S6). Since performance plateaued after 1.5M examples, and thus 1.55M was selected as the optimal training set size. Using this sampling strategy, the final dataset was split into 1.55M training examples, 0.5M validation examples and 0.5 test examples.

### 2.4 Model training

Each model configuration was trained multiple times using independent training runs to account for stochasticity in model initialization and optimization. For each architecture, 10-30 independent training runs were performed and the model with the minimum loss on the “validation” (early stopping) portion of the training set (not the test set) was chosen as the best model. Validation loss across runs is summarized in Suppl. Fig. S10. Training was performed on GPU-enabled nodes provided by the Duke Compute Cluster using NVIDIA A5000 and A6000 GPUs. A single GPU was used for each training run. Most training runs completed in under 15 hours.

### 2.5 Model accuracy calculation

Predictive accuracy was assessed on both quantitative accuracy of effect sizes and binary classification accuracy for variants classified as being functional or non-functional. Quantitative effect size prediction accuracy was evaluated using root mean squared error (RMSE) on held-out STARR-seq test data for both K562 and A549-based models. For classification accuracy, model generalization was assessed using unseen variants from saturation mutagenesis MPRA data reported by Kircher et al [19], which includes measurements across multiple cell types, many of which were different from the cell types on which the model was trained, thus testing the model out-of-distribution.

For comparison, the much larger commercial model AlphaGenome [7] was applied to the same MPRA variant functional/non-functional classification task. Note that unlike BlueSTARR, which was not trained on the test regions, AlphaGenome was trained on the entire genome, including the test regions, possibly giving AlphaGenome a predictive advantage when predicting effects in those same regions, due to data leakage. In addition, AlphaGenome was trained on a much larger variety of cell types than BlueSTARR, possibly providing another predictive advantage on the MPRA data, as each BlueSTARR model was trained on only one cell type. AlphaGenome predictions of RNA-seq and separately for ATAC-seq were used in ROC analyses (Suppl. Fig. S7). Because AlphaGenome cannot always link a noncoding variant to a target gene, predicted RNA-seq values were missing for a large proportion (17,000 / 30,000) of tested variants, and those missing values were replaced with zero values for the ROC analysis.

#### 2.5.1 Accuracy calculation of steady-state predictions

Steady-state STARR-seq activity prediction was evaluated from the reference sequence only using held-out test sequences not seen during model training. For each sequence, the model outputs a predicted quantitative effect size which was compared against the naive estimator for effect size that was calculated from the experimental data. This naive estimator was computed as the ratio of mean RNA counts to mean DNA counts across replicates. Predictive performance was assessed by correlation and the root mean squared error between predicted and observed effect sizes across the test set for each model evaluated.

#### 2.5.2 Accuracy calculation on MPRA variants

MPRA variants were labeled based on experimentally measured regulatory activity following the same criteria described by Kircher et al. [19]. Variants were classified as positive if they exhibited a statistically significant effect (p ≤ 1×10^-5^). Variants were classified as negative if they were not statistically significant (p >1×10^-5^) and had an absolute effect size (|log_2_FC| ≤ 0.05). Only variants with a minimum tag count of 10 (as suggested in Kircher et al.) were included. Variants not meeting these criteria were excluded from analysis.

For each variant, predicted regulatory activity scores were obtained from 2 separate runs of the BlueSTARR model, one for the sequence with the reference allele and one for the sequence with the alternate allele. The regulatory activity score of the variant is the difference between the alternate and reference predictions (since the predictions are in the natural log scale). These activity scores were used to rank the sequences, Model performance was quantified using the area under the receiver operating characteristic curve (AUROC), computed by comparing the predicted scores against the binary labels across all thresholds.

AUROC was calculated separately for each variation of the model separately for all variants, as well as for the enhancer and promoter variants. (Fig. 5, Suppl. Fig. S4).

#### 2.5.3 Accuracy calculation on BIRD allelic variants

To further assess zero-shot generalization, we evaluated BlueSTARR predictions on allelic variants identified by the Bayesian Inference of Regulatory Differences (BIRD) model [20]. Variant-level regulatory activity and significance scores were obtained from BIRD calls for the held-out K562 STARR-seq test data.

For each variant, model predictions were generated independently for the reference and alternate alleles and the predicted allelic effect was defined as the absolute difference between alternate and reference predictions.

Binary labels were assigned based on the reported posterior probability (p-reg) from BIRD. Variants with high confidence regulatory effects (p-reg > 0.9) were labeled as positive while variants with low regulatory evidence (p-reg < 0.3) along with a minimum DNA count (minimum of sum of DNA counts across replicates) threshold greater than 50 were labeled as negative. This additional filtering criteria for the negative class ensures that the variants have sufficient sequencing depth so that their lack of regulatory activity reflects true biological inactivity rather than low read coverage. Variants not meeting these criteria were excluded from the analysis. AUROC was computed by comparing the predicted allelic effect sizes against the binary labels. (Fig. 4A)

### 2.6 Analysis of Constraint

To investigate whether predicted regulatory effects reflect evolutionary constraint, we used trained BlueSTARR models to generate allele-specific predictions across genomic regions representing candidate regulatory elements (open regions) and constitutively closed regions (closed regions), under the hypothesis that variants observed in human populations preferentially occupy regulatory configurations that are less deleterious than alternative nucleotide substitutions. For each genomic position, regulatory activity was predicted for all four possible nucleotides using in silico saturation mutagenesis within a fixed 300 bp sequence context, defined as a sliding window centered on the site of interest. Predictions were then compared within site rather than across sites to reduce sensitivity to sequence context.

To detect enrichment of regulatory configurations among alleles observed in human populations, allele-specific predictions were ranked within each site. At each site, the observed alleles consist of the reference allele and a single-nucleotide variant (SNV) identified in the Genome Aggregation Database (gnomAD), a catalog of genetic variants observed in large human populations relative to the reference genome. The remaining two nucleotides are treated as unobserved alternatives under identical sequence context, allowing comparison between realized human variants and theoretically possible but unobserved substitutions. We then summarized whether observed alleles occupied the highest- or lowest-predicted activity configurations. Sites with ties for the maximum or minimum predicted activity were excluded from these analyses. Under a null model in which observed allele status is independent of predicted activity, observed alleles would occupy extreme ranks with probability 0.5.

Diagnostic analyses revealed systematic nucleotide-specific biases related to GC content in predicted activity, which could confound observed-versus-unobserved comparisons. We therefore implemented a mutation-class–conditioned analysis that compares sites at which a given nucleotide substitution class is the observed reference/SNV pair to background sites at which that same substitution class is not the observed reference/SNV pair. This approach allows inference on regulatory constraint while controlling for nucleotide composition effects and reducing the influence of mutation-dependent biases on enrichment estimates.

To explore whether enrichment patterns vary with genomic proximity to genes, we additionally examined the relationship between enrichment probability and distance to the nearest transcription start site (TSS) in closed regions, motivated by the expectation that regulatory constraint may be stronger in genomic regions proximal to genes. The TSS annotations were obtained from Ensembl BioMart (Ensembl release 110, GRCh38). Sites were stratified into quantile-based distance bins, and the proportion of sites at which an observed allele occupied the maximum predicted regulatory configuration was computed for each bin. A grouped binomial logistic regression using *log*_10_-transformed TSS distance as the predictor was used to assess the presence of a distance-dependent trend. Full methodological details are provided in the Supplementary Methods.

### 2.7 Binding Motif Analysis

To better understand the nature of predicted gain-of-function mutations utilized in the foregoing analysis of constraint, we analyzed allelic changes and their potential impacts on matches to known TF binding motifs. In particular, to identify potential transcription factor binding changes associated with predicted gain-of-function variants, we identified motif occurrences on either DNA strand in both observed sequences containing a variant and dinucleotide-shuffled control sequences. We applied p-value thresholds of p < 10⁻⁵ and z-score > 3 to identify motifs in which binding gains were significantly enriched among the top predicted gain-of-function (GoF) variants. Motif gain and loss events were defined by comparing motif bindings between the two allele-specific sequences. A motif gain event was defined when a motif binding site was significant in the alternative allele sequence but not significant in the observed allele sequence:

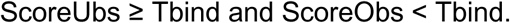

On the other hand, a motif loss event was defined when a motif binding site was significant in the observed allele sequence but not significant in the alternative allele sequence:

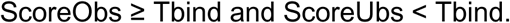

Because multiple motif hits may occur within the sequence window for a given motif, we reduced the results to at most one gain event and one loss event per (variant, motif) pair.

Additional details are provided in Suppl. Text. S1.4.

### 2.8 Predicting transcriptional response to chemical perturbations

In order to evaluate the model’s ability to learn nuanced transcriptional patterns induced by experimental perturbations as reflected in the training data, we sought to reproduce the GR/AP-1 motif spacing experiment described by Vockley et al. [21] that tests the variation in regulatory activity with varying distances between transcription factors GR (glucocorticoid receptor) and AP-1 (a heterodimer consisting of JUN and FOS) in cells treated with a synthetic glucocorticoid drug, dexamethasone (DEX). The original synthetic sequences assayed in that earlier study containing consensus binding motifs for GR and AP1 separated by varying lengths of neutral sequence (20-240 bp) were provided as inputs to the BlueSTARR model to predict activity values (Fig. 10A). Separate BlueSTARR models were trained on STARR-seq data from A549 cells treated with DEX (“A549/DEX”) and A549 cells treated with vehicle control DMSO (“A549/DMSO”). To ensure that the model could capture long-range interactions, we used a 6-layer CNN architecture for this task. The 6-layer architecture provides a larger receptive field than the 5-layer CNN used in all of the earlier analyses. Predicted enhancer activity was measured as the log_2_ ratio between the A549/DEX model and the A549/DMSO model.

## 3. Results

### 3.1 Accuracy of model predictions

#### 3.1.1 Accuracy of steady-state prediction on held-out STARR-seq data

For steady-state prediction of STARR-seq activity from reference sequence only, the tested models all showed a statistically significant correlation (with p = 0.0) in their quantitative prediction of effect size as compared to direct experimental measurements (Fig. 1). The K562 models uniformly performed more accurately than the A549 models, possibly due to differences in insert sizes used in those experiments (mean of 451 bp for K562; 961 bp for A549) (Suppl. Fig. S2). Indeed, in comparing the K562 versus A549 models, the differences are considerably larger than differences between individual K562 models. In particular, steady-state prediction accuracy was highly similar between tested architectures, and between training protocols. A notable exception is the K562 model trained on 1 kbp sequences, which underperformed as compared to the 300 bp models on that cell type, consistent with the distribution of insert sizes from the plasmid libraries used in the K562 experiment having most of its mass at lengths shorter than 1 kbp (Suppl. Fig. S2).

**Figure 1.**
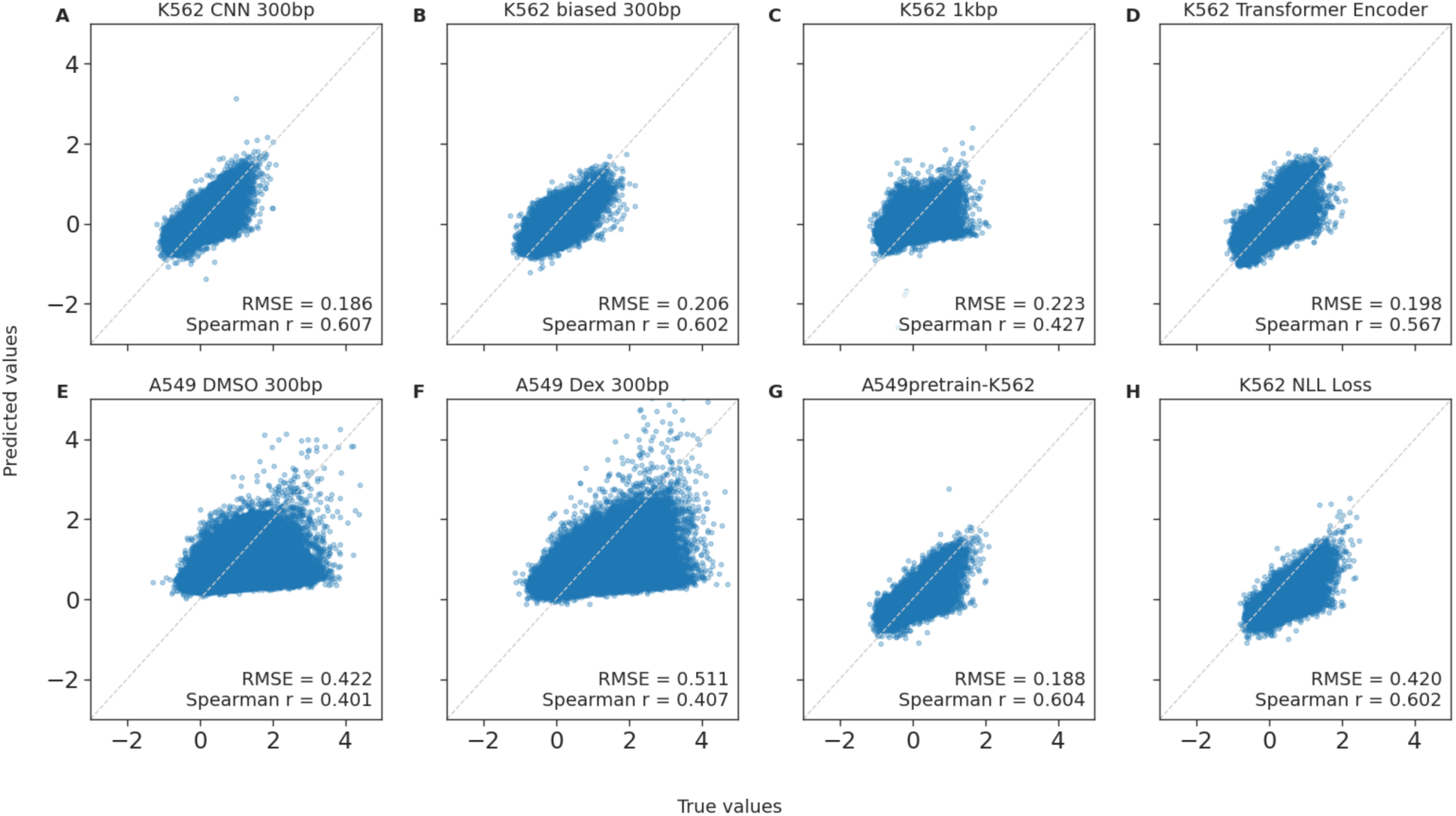
Scatterplots of experimental effect sizes (x-axis) versus predicted effect sizes (y-axis), where effect sizes represent log_2_ ratio of RNA output per unit DNA in reporter assay, and experimental values were computed via ratios of mean read counts across replicates. (A) K562 CNN 300 bp model trained with MSE loss. (B) K562 CNN model trained on data with biased downsampling. (C) K562 model trained on 1 kbp input sequences. (D) K562 model with transformer encoder architecture. (E) A549/DMSO CNN 300bp model trained with MSE loss. (F) A549/DEX CNN 300bp model trained with MSE loss. (G) A549/DMSO CNN model fine-tuned with K562 data trained on 300 bp sequences with a MSE loss function. (H) K562 CNN 300bp model trained with a negative log-likelihood based loss function.

Because each training run in TensorFlow is initialized with random synaptic weights, we have found it necessary to train multiple models of the same architecture and select the model with the best convergence on training data. Partitioning the training data into a training partition and a validation partition (disjoint from the held-out test set), we selected the model with the lowest validation loss and then tested that model on the unseen test set. Performing more training runs generally resulted in the selected model having better generalization to unseen data, with sizable improvements seen from just a few runs (Fig. 2), though minimal improvements were seen when doing more than ten runs (Suppl. Fig. S3). As such, all subsequent results are reported based on the best model of ten training runs, with that best model being selected based on its loss on the validation partition of the training data (not the held-out test set).

**Figure 2.**
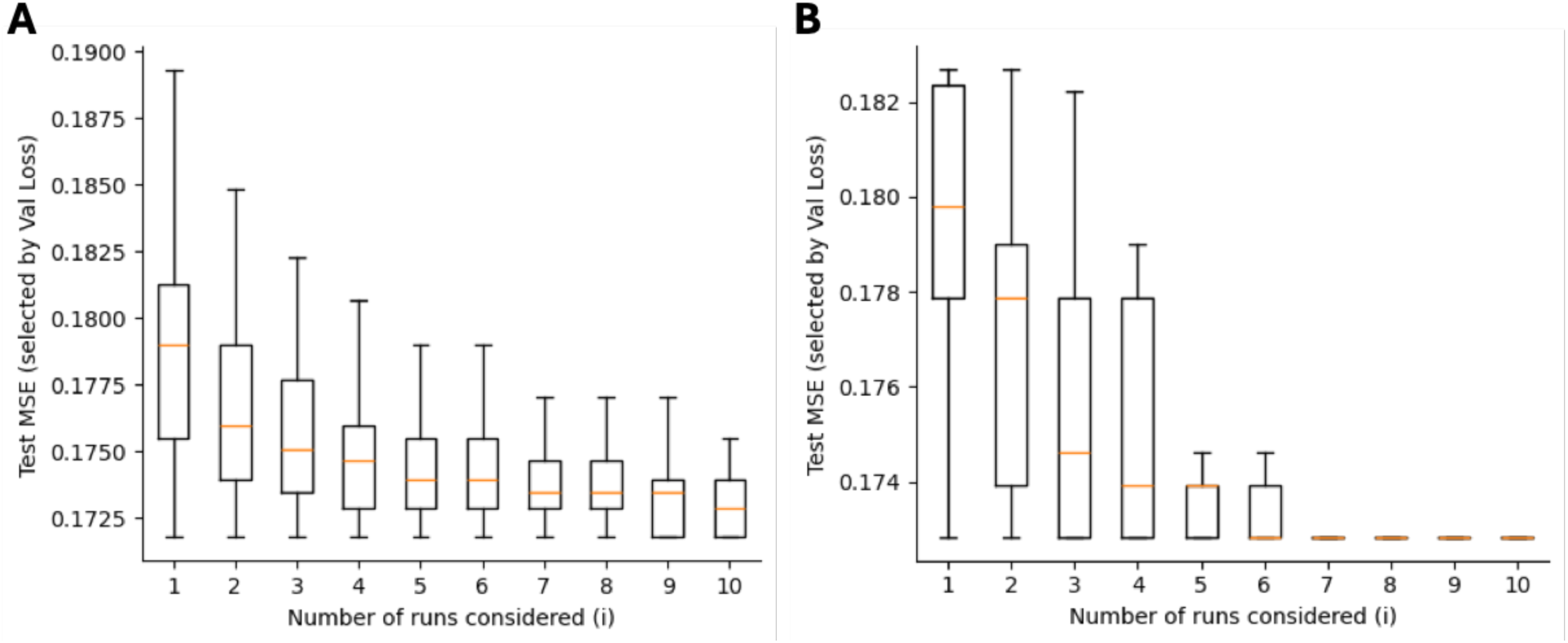
Number of runs (x-axis) required to achieve a given RMSE (y-axis) on simulated data sets. (A) A549/DMSO data for the first 10 runs. (B) K562 data for the 10 runs.

The impact of model architecture and training protocol was found to be relatively minor compared to the impact of the training set, with all of the models training on K562 data performing similarly to each other (Fig. 3). While the 300 bp model with 5 layers produced the lowest RMSE, the 4-layer model mimicking the DeepSTARR architecture performed very similarly, as did the transformer encoder architecture. Even though the biased downsampling methods produce a more uniform distribution of theta across the training set since it over samples underrepresented expression levels, the RMSE of the model trained on biased downsampled data was higher than the model trained on unbiased downsampled data, as did extending input sequence length from 300 bp to 1 kbp. RMSE of the A549 models was substantially higher than that of the K562 models.

**Figure 3.**
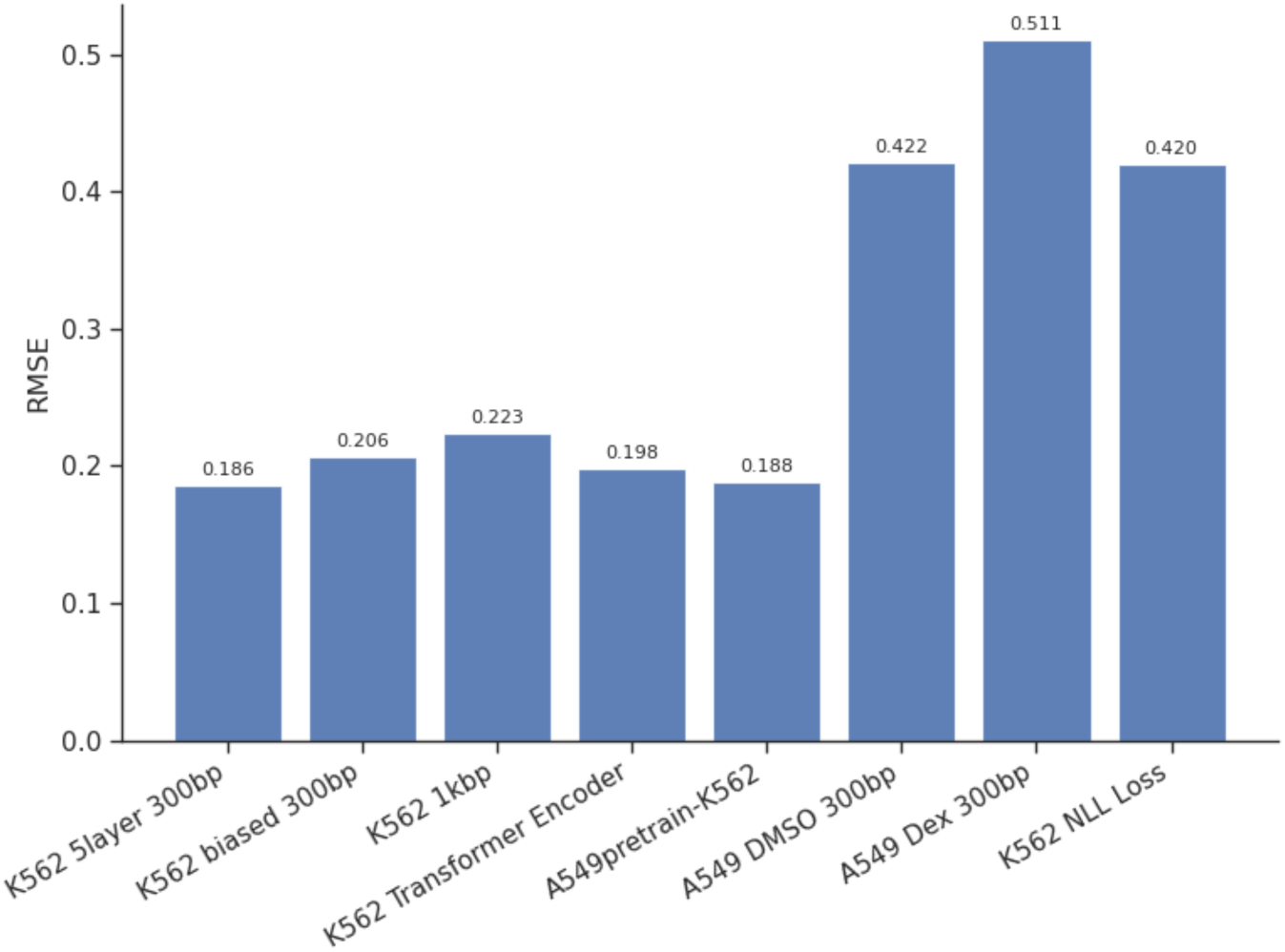
RMSE of steady-state (reference sequence) predictions on held-out STARR-seq data.

#### 3.1.2 Accuracy of zero-shot allelic prediction on held-out STARR-seq and MPRA data

For K562-trained models, AUC (Area Under ROC) values for zero-shot prediction of unseen regulatory variant effects were similar between STARR-seq and MPRA data, with AUCs being comparable on both MPRA and STARR-seq data (Fig. 4A, B). While performance on STARR-seq data yielded an AUC of 0.606, performance on MPRA data increased with the number of convolutional layers, with no additional gain observed beyond 4 layers indicating diminishing returns with increased model depth.

Overall performance is slightly lower on STARR-seq compared to MPRA data. Notably, despite being trained exclusively on K562 cell type data, the model demonstrated robust performance on MPRA dataset, which comprises a diverse set of cell types. This difference may reflect dataset-specific characteristics, including differences in assay design and the diversity of cell types represented in the MPRA dataset (HepG2, HEK293T, HeLa, HaCaT, Neuro-2a, LNCaP, SK-MEL-28, SF7996, K562, HEL92.1.7, NIH/3T3 and Min6) compared to the single-cell type (K562) in the STARR-seq dataset.

**Figure 4.**
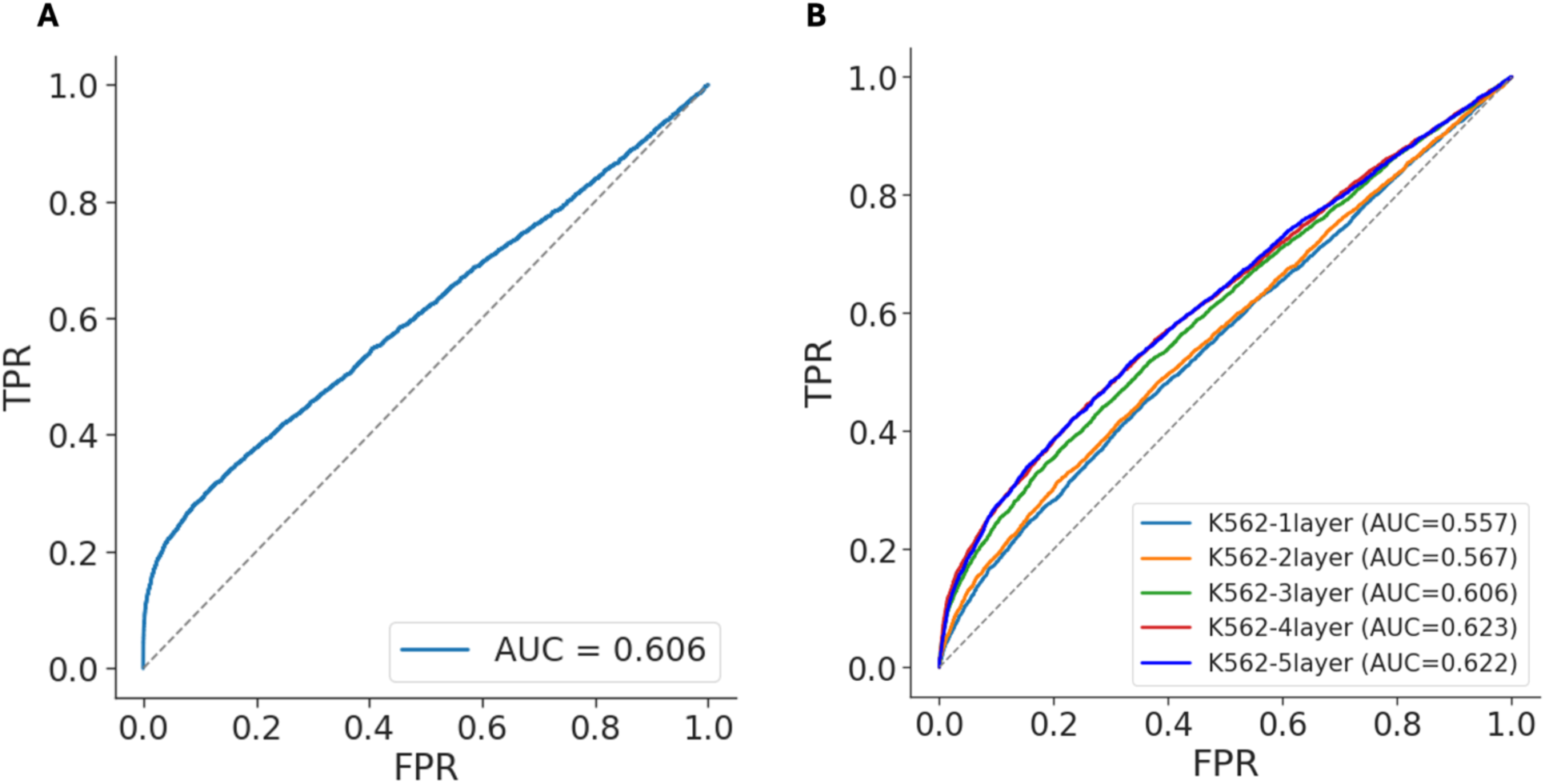
ROC of allelic prediction. (A) ROC for held-out STARR-seq data. (B) ROC for unseen MPRA data, for K562-trained models with one to five convolutional layers.

Trends in MPRA AUC for allelic predictions from models of different architectures (Fig. 5) were largely consistent with trends in steady-state RMSE results described above. Most notably, AUC differed between predictions on enhancers versus promoters. Biased downsampling of K562 training data produced the most accurate convolutional model for promoters, while not producing improvements relative to other models for prediction in enhancers. In contrast, the most accurate model for enhancers was produced via transfer learning by pretraining a model on A549 training data followed by fine-tuning on K562 training data. The negative log likelihood loss function provided no advantage over simple MSE loss. Similarly, the transformer encoder architecture did not produce any advantage over the CNN models.

**Figure 5.**
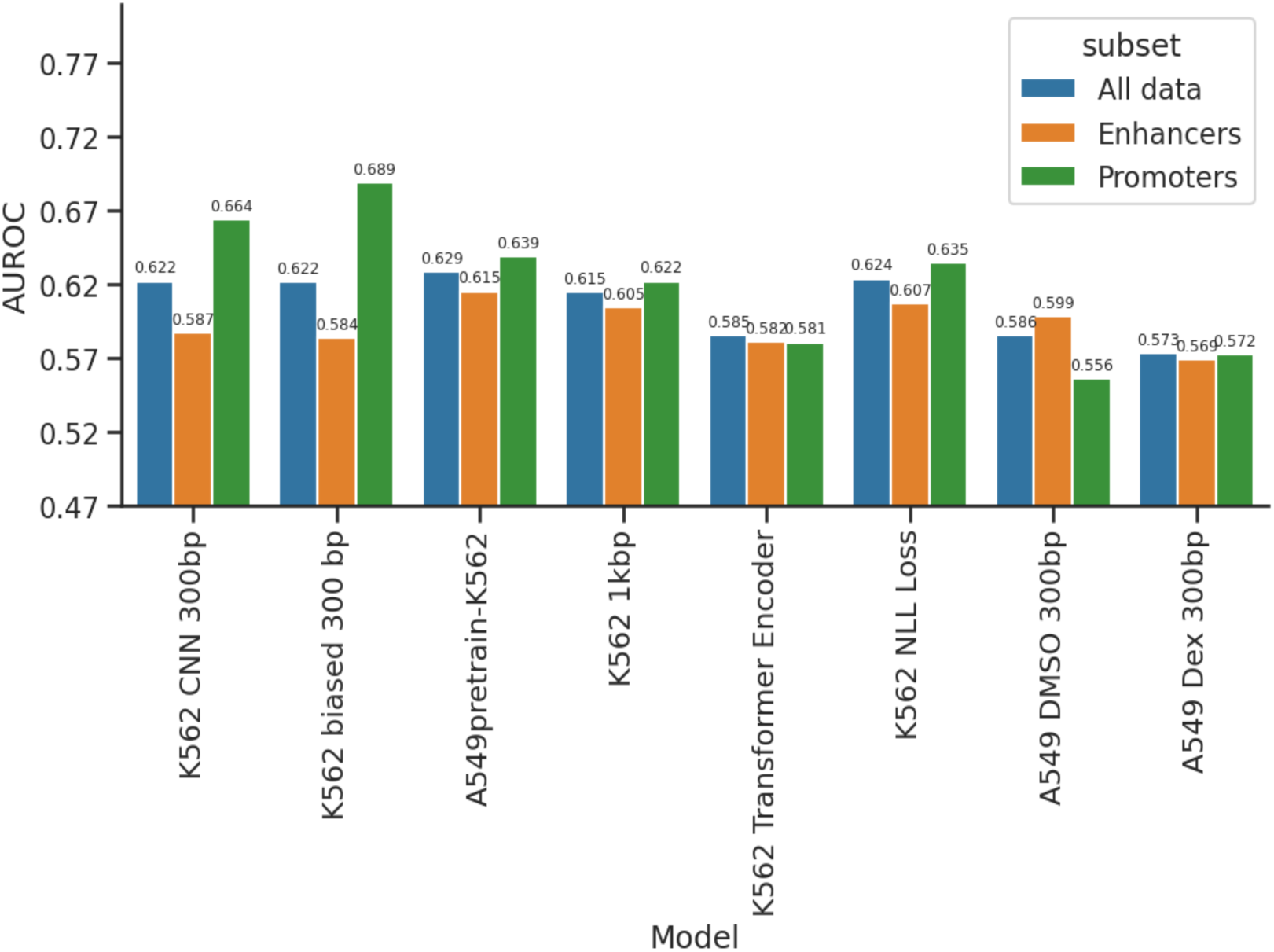
AUC values on MPRA enhancers (orange) and promoters (green) and union set of enhancers and promoters (blue).

AlphaGenome achieved a higher AUC than BlueSTARR on the MPRA data (Suppl. Fig. S7) when the ATAC-seq predictions from AlphaGenome were used, as could be expected given that AlphaGenome was trained on the entire genome, including regions in the test set, possibly leading to data leakage that could inflate measured accuracy. In addition, AlphaGenome was trained on data from a variety of cell types, as compared to BlueSTARR that was trained on a single cell type. AlphaGenome failed to make RNA-seq predictions of 17000/30000 test variants, resulting in lower accuracy when that data modality was used for prediction on MPRA data (Suppl. Fig. S7).

### 3.2 Detecting genome-wide signals of natural selection against gain and loss of regulatory function

Because regulatory activity influences gene expression, variants that substantially alter regulatory output are expected to be subject to purifying selection. When applied on a large scale across the genome, the K562 CNN model revealed significant patterns consistent with natural selection against both loss of function variants in existing cCREs and gain of function variants in constitutively closed (non-cCRE) regions.

#### 3.2.1 Rank-based enrichment of observed alleles

If regulatory activity influences fitness, variants observed in human populations may preferentially occupy regulatory configurations that are less deleterious than alternative nucleotide substitutions. To investigate whether predicted regulatory activity reflects evolutionary constraint, we evaluated whether alleles observed in human populations preferentially occupy extreme predicted regulatory configurations. Using the rank-based framework described in Section 2.6, we estimated the proportion of sites at which an observed allele (reference or single gnomAD

SNV) attains either the minimum predicted activity (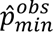) or maximum predicted activity (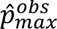). Under the null expectation that observed allele status is independent of predicted regulatory activity, each event should occur with probability 0.5.

##### 3.2.1.1 Closed regions

Across 10,817,580 closed-region sites, the proportion of sites at which an observed allele occupied the minimum predicted activity was 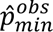 = 0.50295 (95% CI: 0.50265–0.50325), exceeding the null expectation of 0.5 (two-sided *z*-test, *p* = 9.76 × 10^−84^). Conversely, the proportion occupying the maximum predicted activity was 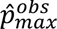 = 0.49417 (95% CI: 0.49387–0.49447), significantly below 0.5 (*p* < 10^−300^).

Although the absolute deviations from 0.5 are small (≈0.3–0.6%), they are consistently estimated across millions of sites. In closed regions, observed alleles are modestly enriched among locally low-activity configurations and depleted among locally high-activity configurations relative to unobserved alleles. Note that as this is a marginalization over millions of sites, it is likely highly diluted by the vast number of regions in which there is likely no effect, thus resulting in effect measurements that appear modest. We expect that the strength of the effect among the subset of sites that are truly under selection to be potentially much stronger.

##### 3.2.1.2 Open regions (K562 cCREs)

Among 2,722,581 open-region sites, the directional pattern was reversed. The proportion of sites at which an observed allele occupied the minimum predicted activity was 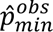 = 0.49766 (95% CI: 0.49707–0.49826), below the null expectation of 0.5 (*p* = 1.30 × 10^−14^). The proportion occupying the maximum predicted activity was 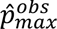 = 0.50462 (95% CI: 0.50402–0.50521), exceeding 0.5 (*p* = 1.92 × 10^−52^).

As in closed regions, the absolute deviations are modest (≈0.2–0.5%) but directionally consistent across millions of sites. In open regulatory regions, observed alleles are modestly enriched among locally high-activity configurations and depleted among locally low-activity configurations (Fig. 6), as expected for putative loss of function regulatory variants.

**Figure 6.**
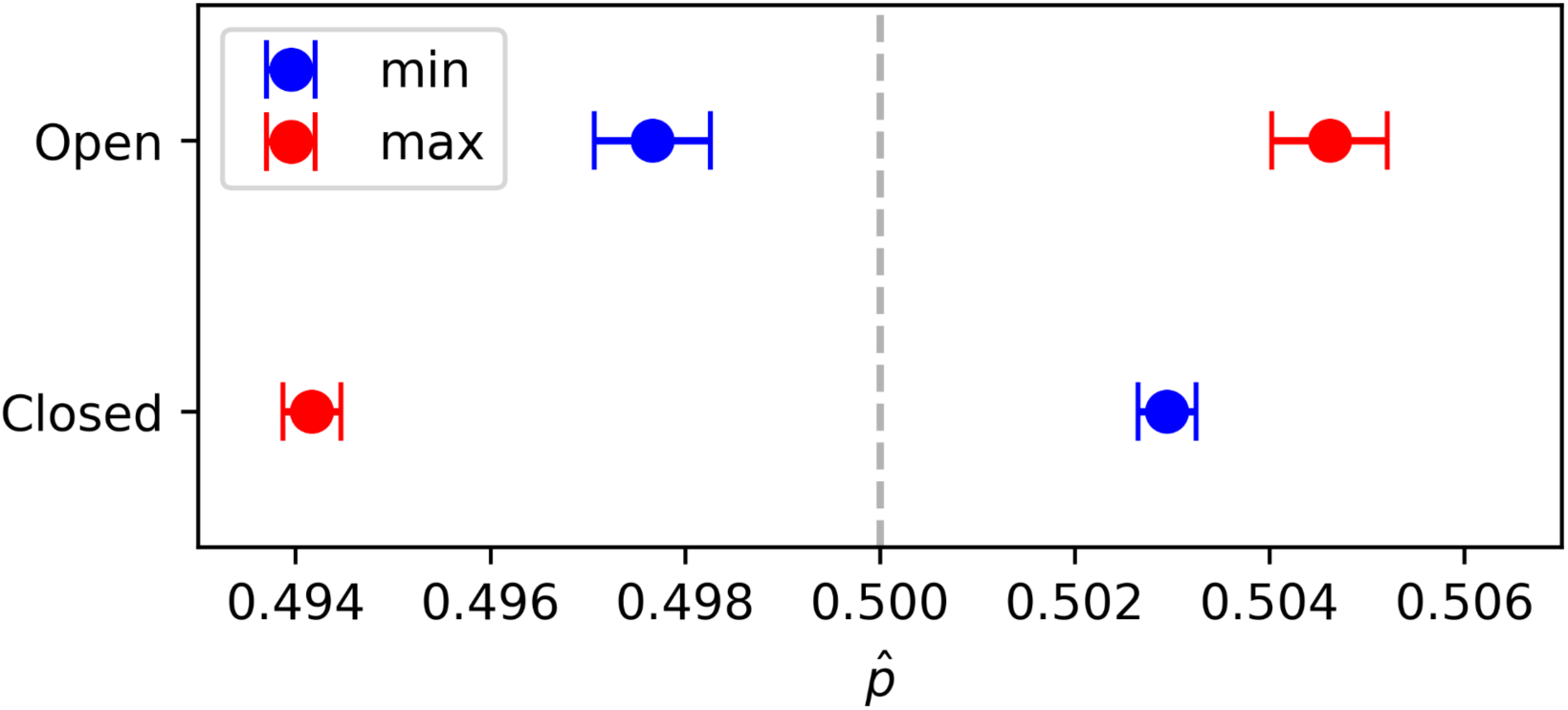
Observed proportions of sites at which an allele present in the human population (reference or gnomAD SNV) occupies the maximum predicted regulatory activity configuration (red) or minimum predicted activity configuration (blue). The dashed reference indicates the null expectation of 0.5. In open regions, observed sites preferentially occupy more active ranks while closed regions exhibit the opposite behavior.

Taken together, these results reveal a context-dependent reversal in the relative occupancy of extreme predicted regulatory states by observed alleles. Observed variants in closed regions tend to occur in locally lower-activity configurations, whereas in open regulatory regions they preferentially occupy higher-activity configurations.

Because the rank-based framework compares observed and unobserved allele groups without conditioning on nucleotide identity (i.e., whether the allele is A, T, C, or G), these enrichment patterns could in principle be influenced by systematic nucleotide-specific biases in the prediction framework or by differences in nucleotide composition across allele classes. We therefore examined whether predicted regulatory activity varies systematically with nucleotide identity and whether nucleotide frequencies differ across variant-type categories.

#### 3.2.2 Rank-based enrichment of observed alleles

To evaluate this possibility, we assessed whether systematic nucleotide-specific biases were present in BlueSTARR predictions and whether nucleotide composition differed across allele categories.

In both closed and open regions, predicted regulatory activity differed significantly across nucleotide identities (Welch’s ANOVA, *p* < 10^−300^). In addition, nucleotide identity was unevenly distributed across variant-type categories (`ref`, `snv`, `unobs`) in both genomic contexts (chi-squared tests of independence, *p* < 10^−300^). Stratified analyses confirmed that nucleotide-specific prediction differences persist within each variant-type class.

These results indicate that both prediction magnitude and nucleotide composition vary systematically with allele identity. Because the primary rank-based enrichment framework does not explicitly condition on nucleotide composition or mutation structure, these structural features could introduce confounding into observed-versus-unobserved comparisons. We therefore implemented mutation-class–conditioned analyses that compare observed and background behavior within matched nucleotide-pair classes.

#### 3.2.3 Mutation-class–conditioned analysis

To determine whether the enrichment patterns observed above persist after conditioning on nucleotide-pair structure, we applied the mutation-class–conditioned framework described in Section 2.6. For each unordered nucleotide pair *p* ∈ {*AT*, *AC*, *AG*, *CG*, *CT*, *GT*}, we compared the within-site contrast statistic *Δ*_*i*_(*p*) between sites at which *p* corresponds to the observed ref/SNV pair and sites at which it does not. Welch two-sample *t*-tests were performed separately for each mutation class within each genomic context, with Bonferroni correction across the six classes.

##### 3.2.3.1 Closed regions

In closed regions, all six mutation classes exhibited negative mean differences (*δ*(*p*) < 0), indicating that when a given nucleotide pair corresponds to the observed ref/SNV configuration, it tends to occur at sites where that pair is relatively lower in predicted regulatory activity compared to its complementary pair. All classes remained significant after Bonferroni correction (Table 1). Although the absolute effect sizes are small (on the order of 10^−4^ − 10^−3^ on the *log*_2_ (*RNA*/*DNA*) scale), the direction of effect is consistent across mutation classes and precisely estimated at this scale.

**Table 1.** Mutation-class-conditioned contrasts in closed regions. *δ*(*p*) = *μ*_*obs*_(*p*) − *μ*_*b*g_(*p*). Bonferroni correction applied across all six classes.

| Class | $N_{Obs}$ | $N_{bg}$ | $\delta(p)$ | $t$ | $p_{Bonf}$ |
| --- | --- | --- | --- | --- | --- |
| AT | 924888 | 9892692 | -7.36e-5 | -3.86 | 6.77e-4 |
| AC | 949999 | 9867581 | -4.36e-4 | -22.86 | 6.79e-115 |
| AG | 3622658 | 7194922 | -8.26e-5 | -6.46 | 6.49e-10 |
| CG | 757579 | 10060001 | -1.03e-5 | -46.92 | <1e-300 |
| CT | 3613387 | 7204193 | -7.04e-5 | -5.49 | 2.38e-7 |
| GT | 949069 | 9868511 | -4.24e-4 | -22.30 | 2.22e-109 |

##### 3.2.3.2 Open regions (K562 cCREs)

In open regions, the mutation-class–conditioned results were more heterogeneous in sign but remained structured (Table 2). For several mutation classes (including AC, AG, CT, and GT), observed pairs were associated with higher relative predicted activity compared to background sites (*δ*(*p*) > 0), whereas for others (notably AT and CG) the opposite pattern was observed.

**Table 2.** Mutation-class-conditioned contrasts in open regions. *δ*(*p*) = *μ*_*obs*_(*p*) − *μ*_*b*g_(*p*). Bonferroni correction applied across all six classes.

| Class | $N_{Obs}$ | $N_{bg}$ | $\delta(p)$ | $t$ | $p_{Bonf}$ |
| --- | --- | --- | --- | --- | --- |
| AT | 142474 | 2580107 | -8.09e-4 | -8.41 | 2.42e-16 |
| AC | 226121 | 2496460 | 1.01e-3 | 13.62 | 1.99e-11 |
| AG | 913672 | 1808909 | 2.08e-3 | 43.57 | <1e-300 |
| CG | 310861 | 2411720 | -3.18e-3 | -46.05 | <1e-300 |
| CT | 903411 | 1819170 | 2.14e-3 | 44.54 | <1e-300 |
| GT | 226042 | 2496539 | 8.70e-4 | 11.57 | 3.52e-30 |

As in closed regions, all mutation classes remained significant after Bonferroni correction, with absolute effect sizes generally on the order of 10^−3^. Despite class-specific heterogeneity, the mutation-conditioned contrasts demonstrate systematic deviations from background expectations.

Taken together, these mutation-class–conditioned analyses indicate that the aggregate enrichment patterns observed in the rank-based framework are not solely attributable to nucleotide-identity imbalance. In closed regions, observed mutation classes consistently occur at sites where those substitutions are predicted to reduce regulatory activity relative to complementary alleles, while in open regulatory regions class-specific contrasts remain detectable after conditioning on mutation type. These findings are consistent with purifying selection acting on both gain- and loss-of-function regulatory variants: predicted activity-increasing variants are depleted in constitutively closed regions, whereas predicted activity-decreasing variants are depleted within active regulatory elements.

#### 3.2.4 Effect of distance on rank enrichment

To explore whether enrichment patterns vary with genomic proximity to genes, we examined the relationship between enrichment probability and distance to the nearest transcription start site (TSS) in closed regions. Sites were stratified into quantile-based distance bins, and the proportion of sites at which an observed allele occupied the maximum predicted regulatory configuration was computed for each bin.

Across bins, enrichment probability exhibited a modest but consistent increase with increasing distance from the nearest TSS (Figure 7). A grouped binomial logistic regression using *log*_10_-transformed TSS distance as the predictor confirmed a significant positive association between enrichment probability and distance (*β* = 0.0113 on the log-odds scale per *log*_10_unit of distance, *p* = 1.08 × 10^−65^).

**Figure 7.**
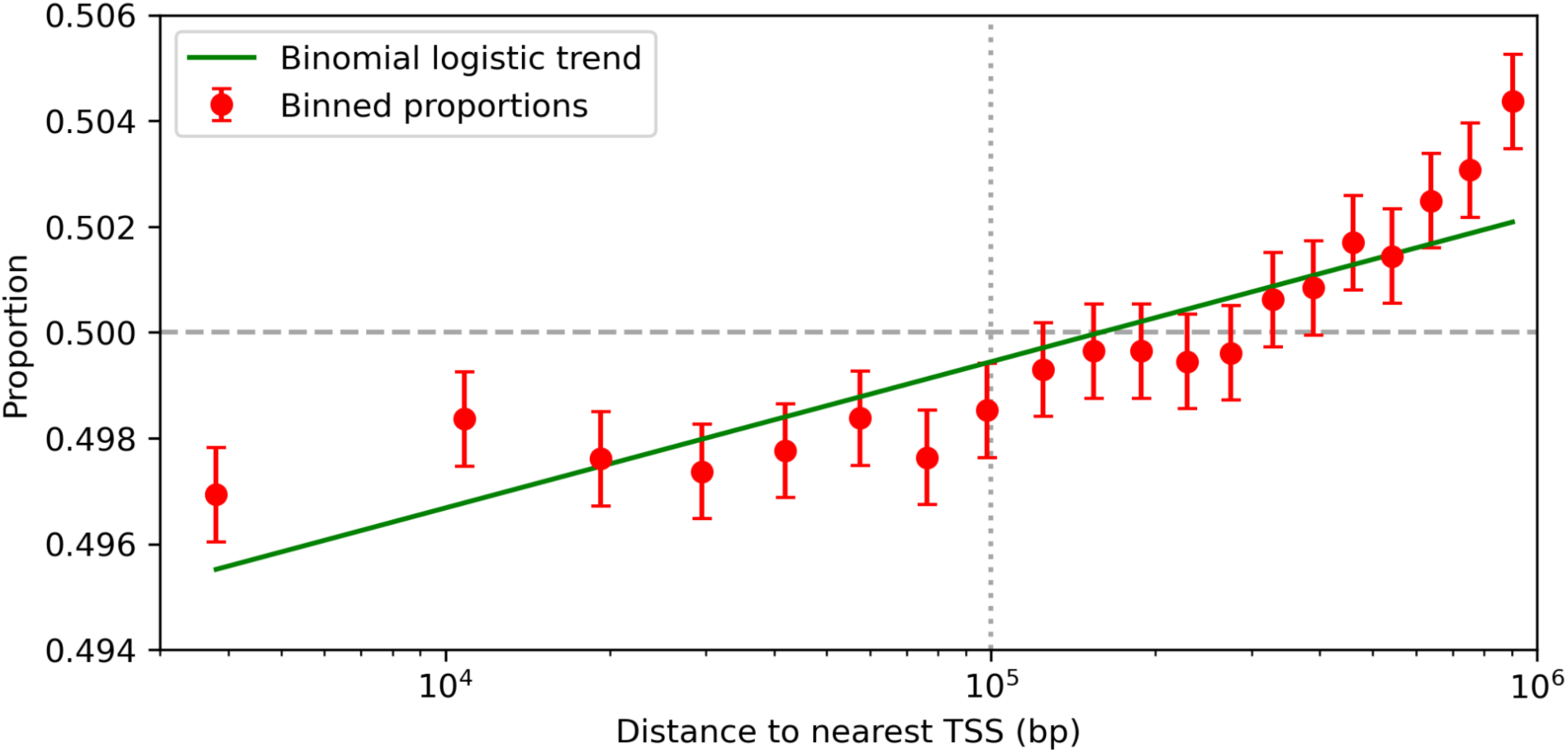
Distance-dependent enrichment of observed alleles in closed regions. Observed proportions of sites at which an allele present in the human population occupies the maximum predicted regulatory configuration are shown for quantile-binned distances to the nearest transcription start site (TSS). Points represent bin-level proportions with 95% confidence intervals. The green curve shows the fitted binomial logistic regression trend using *log*_10_-transformed TSS distance as the predictor. The dashed horizontal line indicates the neutral expectation of 0.5. The dotted vertical line indicates 100Kb from the nearest gene.

Although the magnitude of the effect is small, the trend indicates that observed alleles in closed regions are slightly less likely to occupy locally high-activity configurations at sites proximal to genes. This pattern is consistent with the expectation that regulatory constraint may be strongest in genomic regions near genes.

At finer resolution, however, mutation-direction classes exhibit heterogeneous behavior with respect to distance (Suppl. Fig. S8), indicating that the aggregate trend reflects a mixture of directional mutation effects rather than a single homogeneous mechanism.

#### 3.2.5 Motif analysis

To better understand the nature of the predicted gain-of-function mutations utilized in the foregoing analysis, we analyzed allelic changes and their impacts on matches to known TF motifs (Fig. 8). In particular, to identify potential transcription factor binding changes associated with predicted gain-of-function variants, we identified motif occurrences in both observed sequences containing a variant and dinucleotide-shuffled control sequences. Using p-value thresholds of p < 10⁻⁵ and z-score > 3, we identified 110 motifs (out of 629 tested) in which binding gains were significantly enriched among the top predicted gain-of-function (GoF) variants. Many enriched motifs correspond to known transcriptional activators. Specifically, several members of the bZIP family, including DBP/HLF, CEBPB/CEBPE, ATF/CREB, and FOSL/JUND, were strongly enriched. These factors are known to activate transcription through binding at enhancers and promoters and often recruit coactivators such as CBP and EP300 [22], [23], [24], [25], [26]. In addition, motifs from the ETS family (ELF/ETV) and STAT family (STAT/STAT5A) were also enriched. These transcription factors often act as transcriptional activators, participating in multiple cell signal pathways. [27], [28], [29], [30], [31]. Together, these results indicate that motif gain events enriched among top variants are dominated by transcriptional activators, consistent with the increased regulatory activity predicted by ΔRegulatory Activity.

**Figure 8.**
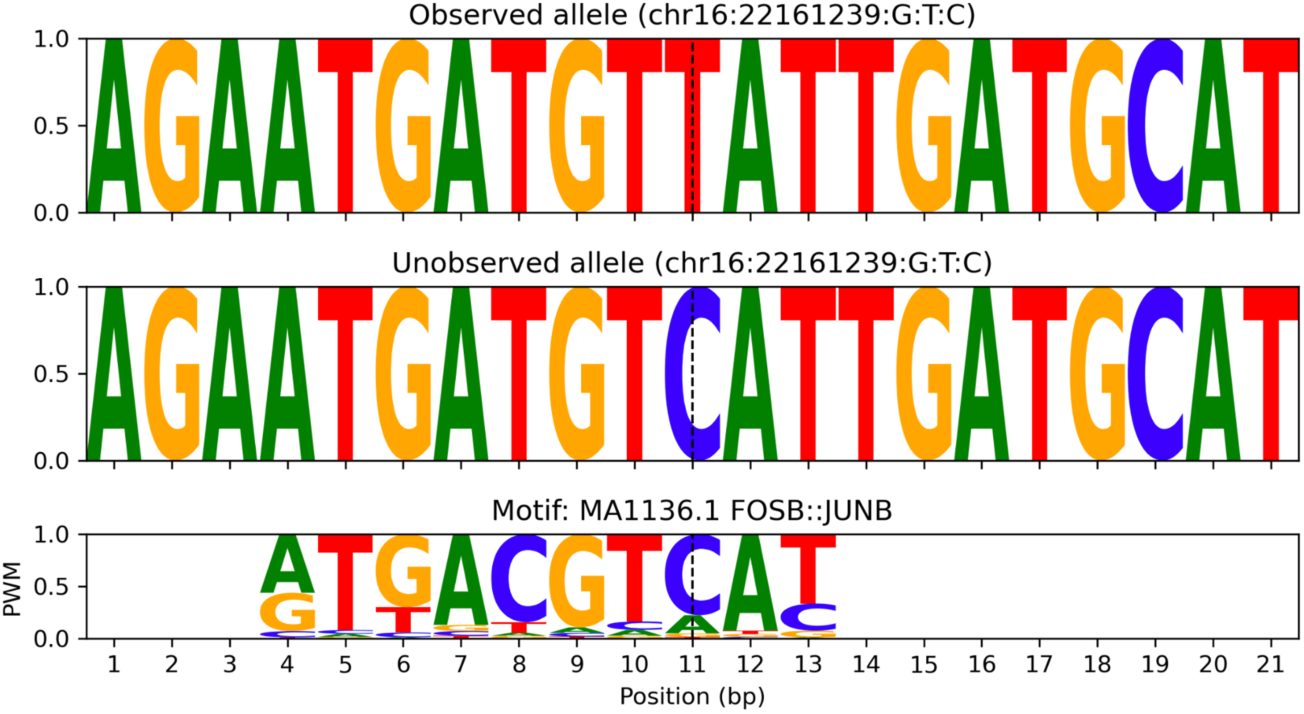
Example mutation from top 1000 ΔRegulatory Activity set that strengthens an AP-1 binding site.

**Figure 9.**
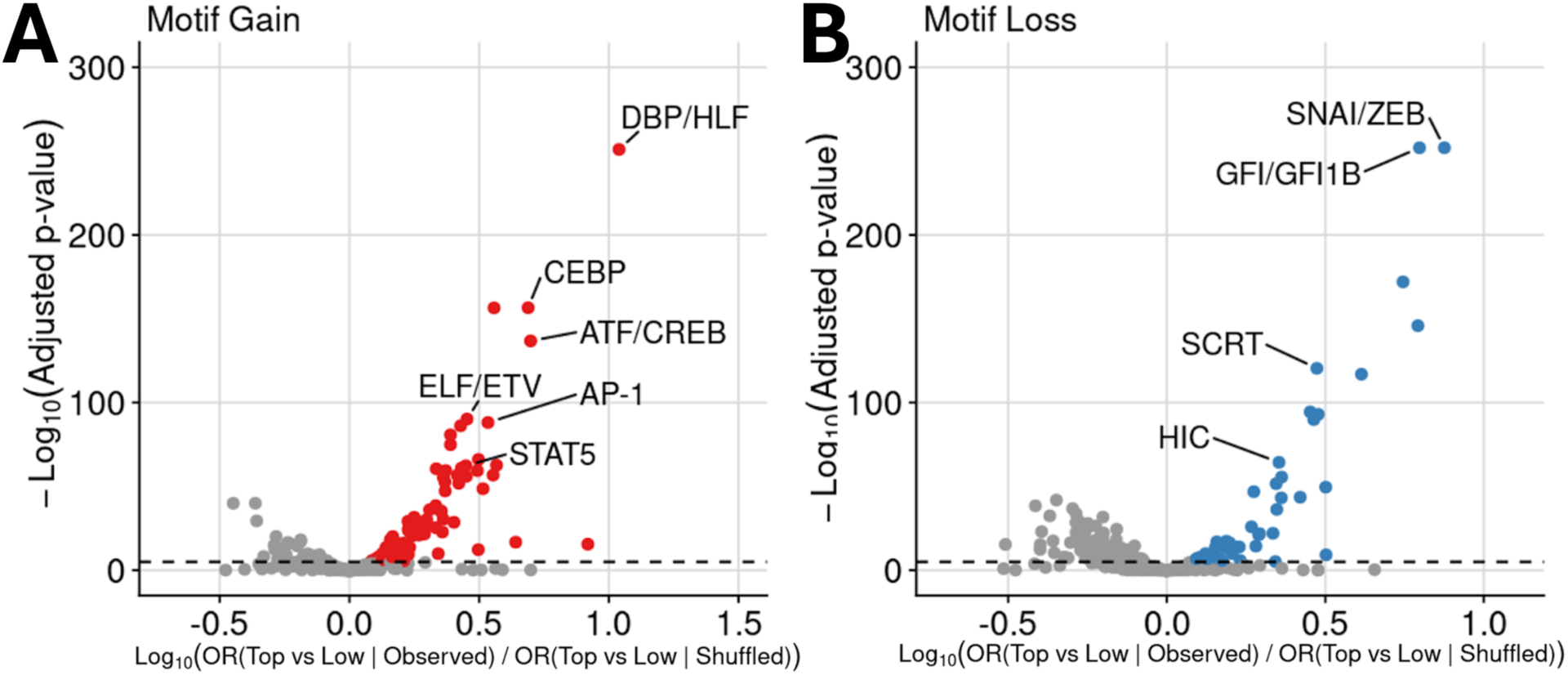
(A) TF motifs enriched for motif gains. (B) TF motifs enriched for motif losses.

In contrast, fewer motif loss events were associated with increased predicted regulatory activity. Using the same thresholds (p-value < 10⁻⁵, z-score > 3), we identified 59 motifs (out of 629 tested) whose binding losses were enriched among the top GoF variants. Many of the enriched motifs belonged to zinc finger transcription factor families, whose regulatory functions are often more diverse and less well characterized than those of classical transcriptional activators. Several zinc-finger families, including ZBTB proteins, can function either as transcriptional repressors or activators depending on cellular context and interacting cofactors [32]. Notably, several well-known transcriptional repressors were identified in our analysis, including SNAI/ZEB, GFI/GFI1B, SCRT, and HIC family factors. These repressors mediate transcriptional repression through recruitment of corepressor complexes. For example, SNAI (Snail) and ZEB transcription factors play key roles in epithelial–mesenchymal transition (EMT) by directly repressing epithelial genes such as CDH1 (E-cadherin) [33], [34], [35].

By controlling for local sequence composition, we demonstrated that these motif binding changes are unlikely to arise from intrinsic sequence bias alone. Instead, they likely reflect biologically meaningful variant regulatory effects consistent with the functional predictions made by BlueSTARR.

### 3.3 Predicting transcriptional responses to chemical perturbations

To evaluate the model’s ability to learn nuanced transcriptional patterns induced by experimental perturbations as reflected in the training data, we sought to reproduce the GR/AP-1 motif spacing experiment described by Vockley et al [21] via model predictions. As shown in Fig. 10B, activity varied as a function of distance between the GR and AP1 motifs, similar to the experimental results in the original publication [21]. Evident in both the experimental and predicted responses are an initial decrease as the transcription factors are moved apart at short distances, then an increase at intermediate distances, another decrease, and then a final increase at long distances that might be indicative of short-range DNA looping in the plasmid [21]. Note that y-axis scales are different from the original publication, likely due to experimental differences, as our model was trained on whole-genome STARR-seq data from another study [14].

**Figure 10.**
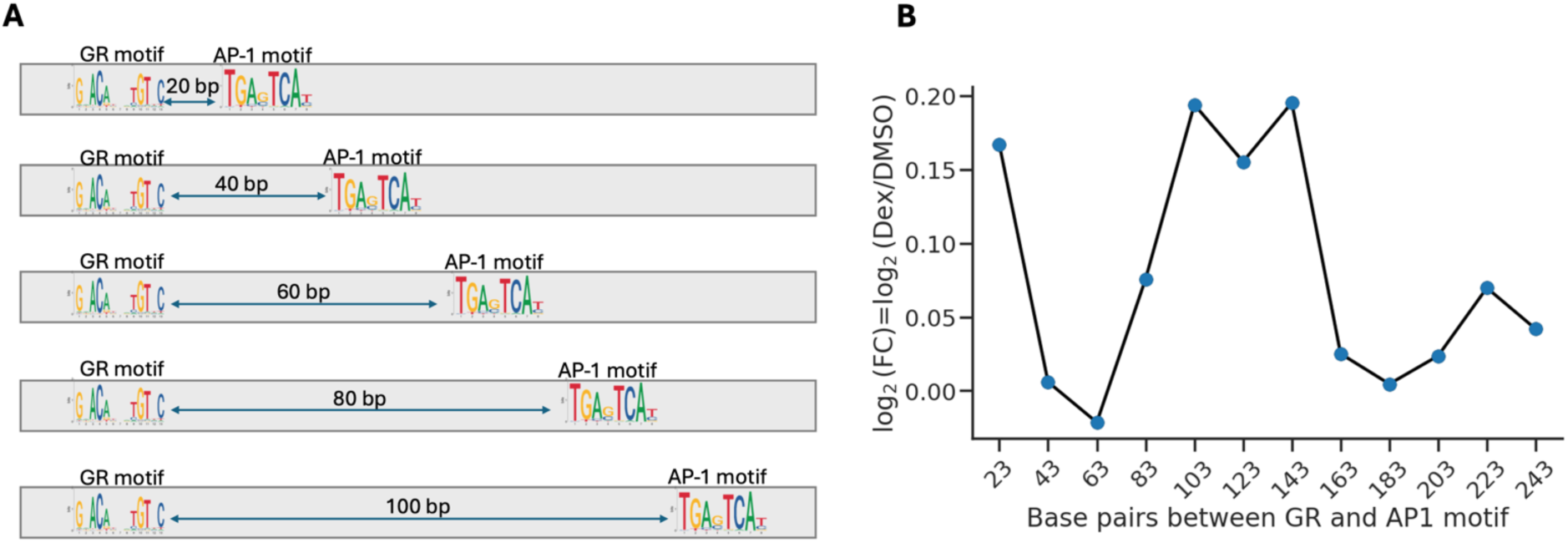
(A) Synthetic sequences tested by Vockley et al. [21] in which a pair of GR and AP-1 consensus motifs were inserted into inert background sequence at a regular progression of lengths. (B) Predicted log2 fold change from BlueSTARR on synthetic sequences containing GR and AP-1 motifs at varying distances.

## 4. Discussion

With the growing appreciation for the influence of the noncoding portion of the genome on human traits, increasing effort has gone into characterizing, either experimentally or *in silico*, the effects of mutations on the cis regulation of genes [2], [3], [4], [5], [7], [8], [9], [10]. While high-throughput reporter assays have now been scaled to test tens of millions of variants at once [12], they remain too laborious to apply in a typical clinical setting, and large-scale experimental studies have to date been able to characterize only a miniscule portion of the space of all possible mutations that could be presented by a patient. Predictive models, on the other hand, can be applied to efficiently impute variant effects at a much larger scale and lower cost than current experimental approaches, but are not as reliable as actual experiments at producing accurate variant annotations, even for the most recent state-of-the-art-methods, as we have shown. Moreover, given the immense space of potential conditions and treatments that may be of interest, a single model may never be available for predicting condition-specific effects for all possible perturbations and cell types in both healthy and disease states.

An opportunity thus exists for the use of combined approaches that utilize unique experimental conditions to generate rich empirical data that can then be used to quickly train lightweight but potentially very specialized predictive models to dissect underlying biology as represented in latent form in the raw experimental data. In such scenarios we view the model less as a general predictor and more as a convenient means of interrogating the latent signals present in the experimental data, and for imputing, under the specific conditions of the experiment, effects of variants that were not directly assayed. While we expect such imputations to have less-than-perfect accuracy, the utility in these models is in their ability to drive rapid hypothesis generation for downstream experimental validation. In this study we described one such framework and demonstrated two use cases, involving drug-dependent gene regulation, and detection of putative signals of natural selection against cis regulatory gain- and loss-of-function mutations. In contrast to industrial-strength models containing billions of parameters that require access to very costly hardware for training, smaller-scale models such as the ones described here can be trained on commodity hardware, in hours rather than weeks or months, and thus can readily serve in aiding interpretation of data from novel experiments.

It has been hypothesized that much of human gene regulation depends on combinatorial binding of multiple transcription factors to CREs, effectively forming a “grammar” that enables fine-scale cellular control of gene expression in different contexts and in response to different stimuli [36], [37], [38]. Whether that grammar enforces a rigid syntax or takes the form of a more flexible “billboard” model [39] has been a subject of some debate, with some recent studies finding evidence in support of both models [40]. In the case of the transcription factor GR, the presence within cells of either endogenous glucocorticoid or synthetic drugs such as dexamethasone can trigger translocation of GR into the nucleus to bind at thousands of sites and alter expression of hundreds of genes [41], [42], [43]. Prior experimental characterization of the relation between GR and its co-factor AP-1 uncovered a pattern of apparent distance-dependent activation in STARR-seq, and prompted the hypothesis that direct GR binding motifs may potentiate clusters of GR binding via indirect tethering via AP-1 [21].

The *in silico* experiments described here suggest that a similar hypothesis could potentially have been formulated using a deep-learning model, by systematically using the model as an “oracle” to predict condition-specific activation for hypothetical CRE sequences. Note that while our model was *not* trained on such synthetic sequences containing GR and AP-1 at a regular progression of distances, the whole-genome STARR-seq data on which it was trained apparently contained sufficient examples of these two factors in proximity across the genome to allow the model, when subsequently run in prediction mode on the unseen synthetic sequences, to reconstruct a nonlinear activation pattern qualitatively similar to that obtained in separate experiments using the synthetic sequences. We believe that this ability to rapidly perform in silico experimentation in pursuit of novel hypothesis generation is an important application for modern deep learning approaches in genomics. We further posit that the ability to train such models quickly on novel experimental data will be key to being able to leverage deep learning methods effectively in biological investigations involving rapid experimental iterations, in which experiments and models are mutually informing and are performed iteratively in alternating fashion for more efficient discovery of novel biology.

Whereas the foregoing example involved an environmental perturbation in the form of a drug treatment, measuring the effects of genetic perturbations is another key application of reporter assays, and imputing mutational effects for variants not captured in those assays is an important application of predictive modeling. Although human variants tested in STARR-seq have been found to have a strong bias toward decreasing rather than increasing expression [12], there are a number of documented cases of cis-regulatory mutations that result in gaining or strengthening of binding sites for transcriptional activators, and consequently deleterious overexpression or ectopic expression of genes [44], [45], [46], [47] referred to here as simply cis-regulatory gain of function. Note that our definition also includes cases of disruption of binding sites for transcriptional repressors, resulting in potentially deleterious gene activation.

While the existence and potential pathogenicity of gain of function mutations have been well-characterized in proteins, particularly in cancer and autosomal dominant diseases [48], a comprehensive search for cis-regulatory gain of function mutations has to our knowledge not been carried out in a systematic manner in the human genome. This is likely due to the much larger search space: while most cis-regulatory loss-of-function mutations will likely occur within annotated CREs, we expect that cis-regulatory gain of function mutations could potentially occur in any location capable of interacting with the promoter of a gene. Given the extensive network of chromatin contacts formed by the three-dimensional genome, that space may be large [49]. *In silico* methods are therefore invaluable for performing large-scale screens for such noncoding gain-of-function candidates. While the proportion of true positives among our predicted gains of function is unknown and may be high, the apparent signal of purifying selection against such mutations when tested at the whole-genome scale suggests that this class of potentially deleterious mutations is worthy of more systematic study.

Indeed, while there is much evidence for the existence of multiple buffering mechanisms against cis-regulatory loss of function mutations, including for example shadow enhancers[50] and redundancy in transcription factors and their binding sites[51], we hypothesize that there may be much less buffering for cis-regulatory gain-of-function mutations, due to the nature of gain-of-function mechanisms, potentially resulting in different selective pressures against such mutations. Additional work is needed to improve on our method for predicting specific cis-regulatory gains of function. Given that such mutations could conceivably act in a condition-specific manner, we posit that there is again a need for easily retrainable models that can predict such mutation effects under very specific conditions as assayed by custom experiments and thus not adequately captured by existing state-of-the-art predictors trained on generic, publicly available data.

Several additional future directions are suggested by the foregoing results. First, while recent state-of-the-art variant effect predictors are prohibitively costly to retrain from scratch on novel experimental data without specialized hardware, due to their massive parameter spaces, fine-tuning of those models [52], perhaps after distillation into smaller parameter spaces [53], [54], may be a practical path toward making use of those “heavyweight” predictors while allowing them to adaptively learn from novel experimental data. One potential challenge for such an approach will be to avoid so-called “catastrophic forgetting” [55], in which the pre-trained model is unable to completely retain its previous knowledge while being re-fit to new data. Thus, refined training methodologies may be needed to allow such fine-tuning approaches to be routinely used for adapting existing large-scale models to novel experimental data. Nevertheless, the result of retraining our A549 model for prediction in K562 cell lines is an encouraging, albeit much smaller-scale, illustration of such a fine-tuning approach.

Second, while methods have recently been developed for generative design of synthetic CRE sequences [11], [56], [57] the need to perform such generative design for specific contexts such as drug responses suggests an additional use case for lightweight models that can be readily trained on novel experimental data. While the models described here are not innately generative, molecule design frameworks employing a *generate-and-test* paradigm have recently been described that include generic models for generating novel sequences that can then be tested via an external “oracle” model [11], [58]. Custom models such as those described here could potentially be used in such scenarios as the oracle.

## Acknowledgements

Research reported in this publication was supported by the National Institute of General Medical Sciences (NIGMS) of the National Institutes of Health (NIH) under award number 1R35-GM150404 to W.H.M., and National Human Genome Research Institute (NHGRI) of NIH 5U01-HG011967 to A.S.A. and W.H.M., and NHGRI 5UM1-HG012053 to T.E.R. Content is solely the responsibility of the authors.

## Data Availability

All the data used for model training are publicly available. The input data used for model training can be downloaded from ENCODE and the accession IDs are given below:

K562 STARR-seq:

Alignment BAM files for DNA library: ENCFF127WNH, ENCFF268POL, ENCFF898TTN

Alignment BAM files for RNA library: ENCFF390VZX, ENCFF405KDE, ENCFF678CBN

A549 STARR-seq:

Alignment BAM files for DNA library: ENCFF978WCI, ENCFF319XHX, ENCFF952VKS, ENCFF351MGI, ENCFF406ZMY

Alignment BAM file for DMSO RNA library: ENCFF616HOR, ENCFF539QGC, ENCFF757DRC, ENCFF708HMK

Alignment BAM files for Dex RNA library: ENCFF604TIK, ENCFF598PMN, ENCFF920RJN, ENCFF820DGO

## Software Availability and Implementation

The code used to performing pre-processing, generate the input data for model training and develop the model are publicly available and is deposited at: https://github.com/Duke-IGVF/BlueSTARR. Pretrained model with the model weights are available for download from: https://huggingface.co/majoroslab/BlueSTARR.

## Author contributions

Model and software development: AT, YC, RV, WHM, HL, BL

Analysis, benchmarking, data preparation: YC, AT, RD, RV, HL, AB, WHM, K-Y K, KD, BL, YD

Writing of manuscript: RV, RD, HL, WHM

Conceptualization and supervision of the work: WHM, ASA, TER

