## Supplementary text and figures for "Modeling gene regulatory perturbations via deep learning from high-throughput reporter assays"

### Supplement for Manuscript “Prioritizing regulatory gain and loss of function variants via deep learning from reporter assays”

#### S1. Supplementary Methods

##### S1.1 Software Implementation

The BlueSTARR command-line tool for training a model accepts three positional parameters: the configuration file; the directory containing training, validation and test data; and the filename stem (which may include a path) for saving the trained Tensorflow model weights and the serialized Keras model object (in the formats used by Keras, which are HDF5 and JSON, respectively). An additional named parameter (—pretrained) can be used to initialize a model from pretrained model weights before training rather than initializing with random values.

The architecture, layout, and hyperparameters for the model to build and train are specified in a configuration file that the BlueSTARR command-line tool reads as one of its required command-line arguments. The configuration file follows a simple text file format in the form of parameter_name = value. Configuration parameters include the number of convolutional layers; number and sizes of convolutional kernels for each layer; drop-out rates (if any); number of transformer encoder layers with their hyperparameters; number of linear layers preceding output layers; the number of epochs to train; and how many epochs to wait before aborting further training after the validation loss is no longer decreasing.

The input format for model training is a FASTA-formatted file of genomic sequence chunks (“bins”) of equal length, and the training, validation, and test count data files. The data files follow a tab-delimited text format, with read counts for each DNA replicate followed by read counts for each RNA replicate. Optionally, the read count columns can be followed by columns denoting the library size to be used for library size-based normalization.

To make reuse of BlueSTARR both easier and also more reproducible and less subject to the specifics of a particular computational environment, we package the entire BlueSTARR suite of tools and related scripts in the form of a Docker container image prebuilt with all dependencies. The Docker container image is available through the GitHub package registry and gets automatically rebuilt every time there is a non-trivial change (i.e., excluding README and other documentation changes) to the main branch of the codebase. Furthermore, we include all Python dependencies in the standard requirements.txt format, which can be used to build a conda environment, and which the Docker container image process uses as well.

#### S1.2 Model architectures

Different versions of the BlueSTARR architecture were evaluated.

##### S1.2.1 Six-layer CNN model

For the A549/DEX and A549/DMSO models used for the predicting transcriptional responses to chemical responses (described in section 3.3 of the paper), a 6-layer convolutional neural network (CNN) model was used to ensure that long range interactions are captured by the model. The 6-layer architecture provides a larger receptive field than the 5-layer CNN model. The rest of the architecture remains the same as the default model described in section 2.2 of the paper.

##### S1.2.2 Transformer Encoder model

To extend the default BlueSTARR architecture, we introduced Transformer encoder layers after the final convolutional layer to enable modeling of long-range dependencies across the input sequence. Specifically the convolutional feature maps were first augmented with rotary positional embedding after which one transformer encoder block was added. Each block consists of layer normalization followed by multi-head self-attention and a position-wise feedforward network, implemented using the Keras *TransformerEncoder* layer. Dropout was applied within each block and a residual skip connection was added around the attention module that could be used.

All transformer-related hyperparameters including the number of layers, number of attention heads, internal dimensionality, and whether to use the residual skip connection were specified through the configuration file.

##### S1.2.3 BlueSTARR architecture with DeepSTARR configuration

This model was trained using the BlueSTARR framework with a DeepSTARR [1] derived configuration on the K562 STARR-seq data. The architecture consisted of four one-dimensional convolutional layers with 246, 60, 60 and 120 filters and kernel sizes of 7, 3, 5 and 3 respectively. All layers had the same padding, dilation factor of 1 and max pooling with size 2. The convolutional layers were followed by flattening and 2 dense layers. No attention layers or global pooling were used.

Training was performed for 200 epochs with a batch size of 128 using Adam optimizer, learning rate of 0.002, dropout of 0.4 in the dense layers and an early stopping “patience” value of 10. Models were trained on sequences of length 300 bp using the same number of training sequences as in the default BlueSTARR model.

##### S1.2.4 Model with finetuning

In this version of the BlueSTARR architecture, models were fine-tuned from pretrained BlueSTARR weights by providing a previously saved weight file (.h5 file) of the A549/DMSO model via the “--pretrained” argument in the script. These pretrained weights were loaded into the network before training the model on the K562 STARR-seq dataset. The rest of the model architecture remains the same as the default BlueSTARR architecture.

#### S1.2.5 Model with input sequence length of 1 kbp

To capture longer-range interactions, we trained the BlueSTARR model using 1000 bp input sequences instead of 300 bp. Data preprocessing was identical to the default setting except that sequences were generated using 1000 bp windows with a 50 bp overlapping step size, in place of the 300 bp windows. All other architectural components and training hyperparameters were unchanged.

#### S1.2.6 Model with Negative Likelihood Loss Function

In this version of the BlueSTARR architecture, we used a negative log-likelihood based loss function instead of the default MSE loss function. Here we derive a *shifted posterior predictive Poisson-gamma mixture* likelihood function for use in training the neural network. Our training objective is to maximize the conditional likelihood of the RNA read counts $Y_j$ for for RNA replicate $j$, conditional on the sum of DNA read counts $X_i$ accross all DNA replicates, and also conditional on the predicted effect size $\hat{\theta}$ produced by the neural network during training, as well as two hyperparameters $\alpha$ and $\beta$ that we shall introduce momentarily:

$$P\left(Y_j \,\middle|\, \sum\nolimits_i X_i, \hat{\theta}, \alpha, \beta\right) \tag{1}$$

As we show later, maximizing this conditional likelihood of the RNA counts given the DNA counts is equivalent to maximizing the joint likelihood of the RNA and DNA counts (See Note #2 below). The total likelihood for an individual training example will be the product of these per-replicate likelihoods under the assumption of conditional independence of the observations given the model parameters. Thus the loss function for one training example will be the negative log of the total likelihood, which will be automatically aggregated across batches by backpropagation during training:

$$Loss = -log \prod_j P\left(Y_j \,\middle|\, \sum\nolimits_i X_i, \hat{\theta}, \alpha, \beta\right) \tag{2}$$

Using this loss function in backpropagation will therefore perform maximum likelihood training of the neural network.

To derive eq. (1), we start with a Poisson likelihood for the DNA counts, parameterized by an expectation parameter $\lambda_X$ that is a random variable with a gamma prior:

$$\sum\nolimits_j X_j \sim Poi(\lambda_X) \tag{3}$$

$$\lambda_X \sim gam(\alpha, \beta) \tag{4}$$

It is well known that a Poisson-gamma mixture is identical to a negative binomial distribution, and is a commonly-used distribution family for sequencing read count data. Because we have no prior expectations as to the DNA count for a randomly-selected STARR-seq insert, we effect a non-informative prior by choosing $\alpha = \beta = 1 \times 10^{-10}$; thus, the observed count $X_i$ will almost entirely dictate the posterior distribution of $\lambda_X$, which we will derive below (see Note #1). The above formulas can be illustrated graphically as:

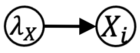


Next we introduce the predicted effect size $\hat{\theta}$ that will serve as a multiplier for $\lambda_X$. This will rescale the Poisson expectation parameter from $\lambda_X$ to $\lambda_Y$, for use in the Poisson likelihood for $Y_j$:

$$Y_j \sim Poi(\lambda_Y) \tag{5}$$
$$\lambda_Y = \hat{\theta}\lambda_X \tag{6}$$

In this way, the predicted $\hat{\theta}$ will represent the relative amount of RNA produced per unit DNA for the current STARR-seq insert. An updated graphical representation is:

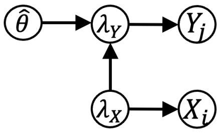


We now make one additional change to this model by incorporating a multiplicative term $L = \frac{RnaLibSize}{DnaLibSize}$ to normalize the Poisson expectations by the ratio of library sizes for DNA and RNA, which we refer to as an *offset*:

$$\lambda_Y = \frac{RnaLibSize}{DnaLibSize}\hat{\theta}\lambda_X \tag{7}$$

Because RNA replicates are generally *biological* replicates via independent transfections, but DNA replicates are typically only *technical* replicates and not necessarily paired with individual RNA replicates, we will replace $X_i$ in the foregoing with simply the sum of counts $X = \sum_i X_i$ across the DNA replicates. Similarly, while $RnaLibSize$ will denote the library size of a particular RNA replicate $j$, the $DnaLibSize$ will denote the sum of library sizes for all DNA replicates.

We can now proceed to derive the conditional likelihood denoted in eq. (1). From the above it should be apparent that this will take the form of a shifted posterior predictive distribution, since we assume that $Y_j$ is drawn from a similar distribution to that of $X$ but with a mean shifted (via multiplication) by the product of $\hat{\theta}$ and the offset $L$.

We begin by noting that if we omit $\hat{\theta}$ and the offset term, then $\lambda = \lambda_X = \lambda_Y$ and $Y_j$ and $X$ would be *iid* under the above model, so that the conditional distribution could be written as the following Poisson-gamma mixture:

$$P(Y_j|X) = \int_0^{\infty} P(Y_j, \lambda|X) d\lambda = \int_0^{\infty} P(Y_j|\lambda) p(\lambda|X) d\lambda \tag{8}$$

due to the conditional independence of $Y_j$ and $X$ given $\lambda$. It can be shown (see Note #1 below) that the posterior $p(\lambda|X)$ for $\lambda_X$ is gamma distributed with parameters given by:

$$\lambda_X|X, \alpha, \beta, n \sim gam(X + \alpha, \beta + n) \tag{9}$$

where $n$ is the number of DNA replicates, which is 1 in our case since we take the sum of counts across DNA replicates as a single observation. Thus we can rewrite eq (8) as:

$$P(Y_j|X, \alpha, \beta, n) = \int_0^{\infty} \frac{\lambda^{Y_j} e^{-\lambda}}{Y_j!} \frac{(\beta + n)^{X+\alpha}}{\Gamma(X + \alpha)} \lambda^{X+\alpha-1} e^{-\lambda(\beta+n)} d\lambda \tag{10}$$

Reintroducing $\hat{\theta}$ and the offset $L = \frac{RnaLibSize}{DnaLibSize}$ into the Poisson likelihood term (and noting that a Jacobian correction is unnecessary as it will ultimately be canceled):

$$P(Y_j|X, \hat{\theta}, \alpha, \beta, L, n) = \int_0^{\infty} \frac{(\hat{\theta} L \lambda)^{Y_j} e^{-\hat{\theta} L \lambda}}{Y_j!} \frac{(\beta + n)^{X+\alpha}}{\Gamma(X + \alpha)} \lambda^{X+\alpha-1} e^{-\lambda(\beta+n)} d\lambda \tag{11}$$

This integral can be solved by substitution, as follows. First, we factor constant terms out of the integral:

$$\frac{(\beta + n)^{X+\alpha} (\hat{\theta} L)^{Y_j}}{\Gamma(X + \alpha)\Gamma(Y_j + 1)} \int_0^{\infty} \lambda^{Y_j+X+\alpha-1} e^{-\lambda(\hat{\theta} L+\beta+n)} d\lambda \tag{12}$$

Substituting $u = (\hat{\theta} L + \beta + n)\lambda$, we have $\frac{du}{d\lambda} = \hat{\theta} L + \beta + n$, and the integral becomes:

$$\int_0^{\infty} \left( \frac{u}{\hat{\theta} L + \beta + n} \right)^{Y_j+X+\alpha-1} e^{-u} \frac{du}{\hat{\theta} L + \beta + n} \tag{13}$$

$$= \frac{1}{(\hat{\theta} L + \beta + n)^{Y_j+X+\alpha}} \int_0^{\infty} u^{Y_j+X+\alpha-1} e^{-u} du \tag{14}$$

Now the integral is in the standard form of a *gamma integral*, which is known to have this solution:

$$\int_0^{\infty} u^{Y_j+X+\alpha-1} e^{-u} du = \Gamma(Y_j + X + \alpha) \tag{15}$$

Reintroducing constant terms that were factored out, the solution to eq (11) can now be written:

$$P\left(Y_j \middle| X, \hat{\theta}, \alpha, \beta, L, n\right) = \frac{(\beta+n)^{X+\alpha}\left(\hat{\theta}L\right)^{Y_j}\Gamma\left(Y_j+X+\alpha\right)}{\Gamma(X+\alpha)\Gamma\left(Y_j+1\right)\left(\hat{\theta}L+\beta+n\right)^{Y_j+X+\alpha}} \tag{16}$$

To see that the above is a negative binomial distribution, choose $p = \frac{\beta+n}{\beta+\hat{\theta}L+n}$ and $r = X+\alpha$ and substitute into the mass function for the negative binomial:

$$P\left(Y_j \middle| r, p\right) = \binom{Y_j+X+\alpha-1}{Y_j}\left(\frac{\hat{\theta}L}{\beta+\hat{\theta}L+n}\right)^{Y_j}\left(\frac{\beta+n}{\beta+\hat{\theta}L+n}\right)^{X+\alpha} \tag{17}$$

$$= \frac{\Gamma\left(Y_j+X+\alpha\right)\left(\hat{\theta}L\right)^{Y_j}(\beta+n)^{X+\alpha}}{\Gamma\left(Y_j+1\right)\Gamma(X+\alpha)\left(\beta+\hat{\theta}L+n\right)^{Y_j+X+\alpha}} \tag{18}$$

which is identical to the r.h.s. of eq. (16). Hence, the conditional likelihood in eq. (1) used in our negative log likelihood loss function is a posterior predictive negative binomial where the mean of the RNA counts is shifted by the multiplicative factor $\hat{\theta}L$ from that of the summed DNA counts.

**S1.2.6.1 Note #1: Demonstrating that the posterior of $\lambda_X$ is gamma distributed**

Here we show that if we have $n$ observations $X_i \sim Poi(\lambda)$ with prior $\lambda \sim gam(\alpha, \beta)$, then the posterior of $\lambda$ is:

$$\lambda | X, \alpha, \beta, n \sim gam(X+\alpha, \beta+n) \tag{19}$$

for $X = \sum_i X_i$. We start by writing the posterior of $\lambda$ via Bayes' rule:

$$p(\lambda|\mathbf{X}, \alpha, \beta) = \frac{p(\lambda, \mathbf{X}|\alpha, \beta)}{P(\mathbf{X}|\alpha, \beta)} = \frac{p(\lambda, \mathbf{X}|\alpha, \beta)}{\int_0^\infty P(\lambda, \mathbf{X}|\alpha, \beta)\, d\lambda} = \frac{P(\mathbf{X}|\lambda)p(\lambda|\alpha, \beta)}{\int_0^\infty P(\mathbf{X}|\lambda)p(\lambda|\alpha, \beta)\, d\lambda} \tag{20}$$

where $\mathbf{X} = \{X_i\}$ is the vector of $n$ observations, and the last step in eq. (20) follows from conditional independence of $\mathbf{X}$ from $\alpha$ and $\beta$ given $\lambda$. Since the $X_i$ are *iid*, the above becomes:

$$p(\lambda|\mathbf{X}, \alpha, \beta) = \frac{p(\lambda|\alpha, \beta)\prod_{i=1}^n P(X_i|\lambda)}{\int_0^\infty p(\lambda|\alpha, \beta)\prod_{i=1}^n P(X_i|\lambda)\, d\lambda} \tag{21}$$

The denominator can be rewritten using the gamma prior and Poisson likelihood:

$$\int_0^\infty \frac{\beta^\alpha \lambda^{\alpha-1}e^{-\lambda\beta}}{\Gamma(\alpha)}\frac{e^{-n\lambda}\lambda^X}{\prod_{i=1}^n X_i!}d\lambda = \frac{\beta^\alpha}{\Gamma(\alpha)\prod_{i=1}^n X_i!}\int_0^\infty e^{-\lambda(n+\beta)}\lambda^{\alpha-1+X}\, d\lambda \tag{22}$$

Substituting $u = \lambda(\beta+n)$ and $\frac{du}{d\lambda} = \beta+n$, the integral in this denominator can be solved via:

$$\int_0^\infty \left(\frac{u}{\beta+n}\right)^{\alpha-1+X} e^{-u}\frac{du}{\beta+n} = \frac{1}{(\beta+n)^{X+\alpha}}\int_0^\infty u^{X+\alpha-1}e^{-u}du = \frac{\Gamma(X+\alpha)}{(\beta+n)^{X+\alpha}} \tag{23}$$

by again making use of a gamma integral on the r.h.s. Thus the complete denominator is:

$$\frac{\beta^\alpha \Gamma(X+\alpha)}{(\beta+n)^{X+\alpha}\Gamma(\alpha)\Gamma(X_i+1)} \tag{24}$$

From the above it can be trivially seen that the numerator is:

$$\frac{e^{-n\lambda}\lambda^X}{\Gamma(X_i+1)}\frac{\beta^\alpha\lambda^{\alpha-1}e^{-\lambda\beta}}{\Gamma(\alpha)} \tag{25}$$

and thus the posterior of $\lambda$ is:

$$p(\lambda|\mathbf{X},\alpha,\beta) = \frac{\frac{e^{-n\lambda}\lambda^X}{\Gamma(X_i+1)}\frac{\beta^\alpha\lambda^{\alpha-1}e^{-\lambda\beta}}{\Gamma(\alpha)}}{\frac{\beta^\alpha\Gamma(X+\alpha)}{(\beta+n)^{X+\alpha}\Gamma(\alpha)\Gamma(X_i+1)}} = \frac{e^{-\lambda(\beta+n)}\lambda^{X+\alpha-1}(\beta+n)^{X+\alpha}}{\Gamma(X+\alpha)} \tag{26}$$

which is $gam(X+\alpha, \beta+n)$.

##### S1.2 Note #2: Maximizing $P(Y|X,\theta)$ with respect to $\theta$ is equivalent to maximizing $P(Y,X|\theta)$

Maximizing the joint likelihood of $Y$ and $X$ with respect to parameter $\theta$ is equivalent to minimizing the negative log of the joint likelihood:

$$\theta^* = \underset{\theta}{argmin} - \log P(Y,X|\theta) = \underset{\theta}{argmin} - \log[P(Y|X,\theta)P(X|\theta)] \tag{27}$$

Given that $X$ is marginally independent of $\theta$ in our model, this is equivalent to:

$$\theta^* = \underset{\theta}{argmin} - \log[P(Y|X,\theta)P(X)] \tag{28}$$

and because $P(X)$ is constant with respect to the argmin and is non-negative, it therefore suffices to minimize the conditional log likelihood of RNA given DNA:

$$\theta^* = \underset{\theta}{argmin} - logP(Y|X,\theta) \tag{29}$$

The default architecture is illustrated in Suppl. Fig. S9.

### S1.3 Analysis of constraint

#### S1.3.1 Genomic interval preparation

This section describes how genomic interval inputs were converted into site-level analytic tables used in downstream statistical analyses. Two genomic region sets were analyzed: a set of constitutively closed ("closed") regions and a set of open regions defined by candidate cis-regulatory elements (cCREs). All analyses were conducted on the hg38/GRCh38 reference assembly.

Closed regions were defined using a previously established exclusion-based pipeline. Briefly, protein-coding genes and regulatory annotations from Ensembl, DNase I hypersensitive regions, and ultra-conserved elements were merged and removed from the genome. Repetitive and technically unreliable sequence, including RepeatMasker annotations, segmental duplications, and assembly gaps, was also excluded. The remaining sequence was restricted to high-coverage regions in gnomAD v3.1, defined as positions with at least 10× coverage in at least 70% of samples. The output of this procedure was a set of high-confidence constitutively closed genomic intervals.

Open regions were defined using candidate cis-regulatory elements (cCREs) from the ENCODE SCREEN database (ENCODE v3) [2]. Because BlueSTARR was trained on K562 reporter assay data, analyses were restricted to cCREs annotated for K562 cells to align the genomic application with the model training context. To promote comparability with the closed set and restrict analyses to regions with reliable variant calls, cCREs were intersected with the gnomAD v3.1 high-coverage mask, and only elements with greater than 80% overlap with high-coverage sequence were retained.

To prevent leakage from model training into evaluation analyses, any region overlapping the BlueSTARR training intervals was excluded. Overlap was defined as any base-pair overlap between a candidate region and a training interval, and exclusions were applied using the true (unpadded) region intervals.

For each retained interval, the nearest transcription start site (TSS) was identified using Ensembl gene annotations (hg38) and bedtools, recording the nearest gene and absolute distance to the nearest TSS.

#### S1.3.2 In silico saturation mutagenesis and prediction generation

Allele-specific regulatory predictions were generated using trained BlueSTARR models, which predict reporter activity from 300 bp genomic sequences. In this work the model is treated as a fixed predictive instrument; the focus of the analysis is on generating comparable allele-level predictions across genomic sites rather than on model training or architecture.

For each genomic region, genomic sequence was extracted from the hg38 reference assembly and evaluated using a fixed-length sliding window of 300 bp. To enable prediction at every base within each interval while maintaining constant input length, regions were padded by extending the interval 150 bp upstream and 149 bp downstream. Predictions were generated for every window position across the padded interval, but downstream analyses retained only predictions centered on bases within the original, unpadded region.

Allele-specific predictions were generated via in silico saturation mutagenesis. At each genomic position, four sequences were constructed by substituting the central base of the 300 bp window with each of the four nucleotides (A, C, G, and T). Each sequence was evaluated independently by the BlueSTARR model, producing four predicted regulatory activity values for that site under alternative nucleotide configurations. BlueSTARR outputs predictions on the natural log scale, ln(RNA/DNA), which were converted to $\log_2$(RNA/DNA) prior to downstream analysis.

Because all four allele predictions at a site share identical sequence context and window placement, they are directly comparable within site. Subsequent analyses therefore focus on relative ordering and contrasts among allele-specific predictions rather than on absolute prediction magnitudes.

##### S1.3.3 Allele annotation and classification

Allele-specific regulatory predictions generated as described above were annotated with population genetic and genomic context information to enable comparisons between alleles observed in human populations and those that are theoretically possible but unobserved.

Population variation data were obtained from gnomAD v3.1. Analyses were restricted to genomic positions at which exactly one single-nucleotide variant (SNV) was observed and no other variant types (e.g., multi-allelic SNVs or indels) were present. At each such site, the reference allele and the single observed SNV allele were classified as observed, while the remaining two nucleotides were classified as unobserved. This yields, for each retained site, a set of four allele-specific predictions partitioned into two observed alleles and two unobserved alleles under identical sequence context.

This restriction ensures a consistent comparison framework across sites, in which the probability that an observed allele occupies any given rank under a null model of independence is 0.5. Throughout the analysis, comparisons are therefore performed between observed and unobserved allele groups within site rather than across sites, reducing sensitivity to differences in baseline predicted activity across genomic regions.

In addition to population annotations, each genomic position was annotated with its distance to the nearest transcription start site (TSS) using Ensembl gene annotations (hg38) and bedtools. TSS distance was retained as a continuous variable and used in both filtering and downstream stratified analyses. For primary analyses, positions more than 100 kb from the nearest TSS were excluded, while analyses explicitly examining distance effects used an extended range up to 1 Mb.

##### S1.3.4 Rank-based enrichment analysis

Allele-specific regulatory predictions yield four predicted activity values at each genomic position, corresponding to the four possible nucleotides evaluated under identical sequence context. We employed a rank-based framework to summarize these predictions in a manner that is robust to model calibration and comparable across sites.

For each site, the four allele-specific predictions were ranked from highest to lowest predicted activity. Analyses were restricted to sites with exactly one gnomAD SNV, yielding two observed alleles (reference and SNV) and two unobserved alleles at each position. The primary outcomes of interest were whether an observed allele occupied the maximum or minimum predicted regulatory configuration within site.

Specifically, we defined two indicator variables:

- $I_{max} = 1$ if an observed allele attains the highest predicted activity at a site, and 0 otherwise;
- $I_{max} = 1$ if an observed allele attains the lowest predicted activity at a site, and 0 otherwise.

Under a null model in which observed allele status is independent of predicted activity, each site contains two observed and two unobserved alleles, implying that the probability an observed allele occupies either extreme rank is 0.5. Enrichment or depletion of observed alleles among extreme predicted configurations was therefore assessed by comparing the observed proportions of $I_{max}$and $I_{min}$ to this null expectation.

Because ranking is performed within site, this framework is invariant to monotonic transformations of the prediction scale and reduces sensitivity to site-specific differences in baseline predicted activity.

#### S1.3.5 Assessment of nucleotide and variant-type bias

To evaluate whether nucleotide identity or mutation structure could influence the enrichment analysis, we assessed nucleotide-specific patterns in predicted regulatory activity and allele composition.

Predicted regulatory activity was compared across nucleotide identities (A, C, G, T) using Welch's one-way analysis of variance (ANOVA), which is robust to unequal variances and sample sizes. Post hoc pairwise comparisons were performed using the Games–Howell procedure. To reduce computational burden while preserving precision, these analyses were conducted on a fixed random subsample of allele-level observations.

To assess whether nucleotide composition differed across allele categories, contingency tables of nucleotide identity by variant-type (`ref`, `snv`, and `unobs`) were constructed and tested for independence using Pearson's chi-squared test.

Finally, to evaluate whether nucleotide-specific prediction differences persisted within allele categories, the above analyses were repeated within each variant-type stratum.

#### S1.3.6 Mutation-class–conditioned analysis

To account for potential confounding due to nucleotide identity and mutation structure, we performed mutation-class–conditioned analyses that compare observed and background behavior within matched nucleotide substitution classes.

Mutation classes were defined as the six unordered nucleotide pairs $\{AT, AC, AG, CG, CT, GT\}$, corresponding to all possible single-nucleotide substitutions up to direction. For each genomic site $i$, allele-specific predictions were available for all four nucleotides, and the observed mutation class was defined by the unordered pair of the reference allele and the single gnomAD SNV at that site.

For each site $i$ and mutation class $p$, we defined a within-site contrast statistic:

$$\Delta_i(p) = \frac{1}{2}(X_{i,p_1} + X_{i,p_2}) - \frac{1}{2}(X_{i,\bar{p}_1} + X_{i,\bar{p}_2})$$

where $\{p_1, p_2\}$ are the nucleotides in class $p$ and $\{\bar{p}_1, \bar{p}_2\}$ are the complementary nucleotides. This statistic compares the mean predicted activity of alleles in class $p$ to that of the complementary pair within the same site.

For each mutation class $p$, sites were partitioned into two groups: those at which $p$ corresponds to the observed reference/SNV pair and those at which it does not. Each site contributes one value $\Delta_i(p)$ to each mutation-class–specific comparison.

For each class, the distributions of $\Delta_i(p)$ were compared between observed and background site sets using Welch's two-sample t-test, allowing for unequal variances and sample sizes. Because six mutation classes were evaluated, p-values were adjusted using Bonferroni correction within each genomic context.

All mutation-class–conditioned analyses were performed separately for closed and open region sets.

#### S1.4 Motif Analysis

To investigate whether genomic sequence mutations can increase regulatory activity by creating or strengthening transcription factor (TF) binding sites, we analyzed the TF binding effects of variants identified using BlueSTARR. For each variant locus, we compared the predicted regulatory activity of the observed allele (reference or population SNP) with that of the corresponding unobserved allele. Variants were evaluated using the change in predicted regulatory activity from the BlueSTARR model (ΔRegulatory Activity = Score(unobserved) − Score(observed)). Variants with positive ΔRegulatory Activity were classified as gain-of-function candidates and used for downstream motif analysis. We identified a subset of variants for which the unobserved allele was predicted to increase regulatory activity relative to the observed allele, suggesting that these mutations may enhance enhancer activity. To explore the mechanistic basis of these predictions, we performed transcription factor motif scanning at variant loci and calculated the change in motif score between alleles (Δmotif).

To compare results across different motifs, the probabilities in each PWM were first converted into log-odds ratios relative to an empirical background model summarized from the flanking regions of those variants. Motif scores were then computed as the log-odds scores, representing the log-likelihood ratio of observing a sequence under the motif model relative to an empirical background model. This score provides a normalized measure of transcription factor binding strength that is comparable across motifs of different lengths. Each sequence was scanned for motif occurrences on both the forward and reverse strands using log-odds motif scoring. For a given motif, the scanning procedure produces a score representing the likelihood of transcription factor binding at each possible position within the sequence window.

To determine whether a motif was considered bound at a given position, motif scores were compared against motif-specific binding thresholds (Tbind). These thresholds represent the minimum score required for a motif instance to be considered a significant binding site. A motif was defined as significantly bound at a position if the motif score exceeded the motif-specific threshold Tbind. For each motif, we estimated the null distribution of log-odds scores under the empirical background using a dynamic programming algorithm that recursively combines per-position motif contributions to obtain the full probability mass function of motif scores under the background model [3]. The binding threshold Tbind was defined as the critical score

corresponding to a right-tail probability of $\alpha = 10^{-3}$, such that $P(S \geq Tbind) \sim 10^{-3}$. Because motif length and information content vary across motifs, the value of Tbind differs for each motif.
The change in motif score between alleles was calculated as

$$\Delta(\text{Motif}) = \text{ScoreUbs} - \text{ScoreObs},$$

where ScoreObs is the motif score for the sequence containing the observed allele and ScoreUbs is the motif score for the sequence containing the alternative allele. This Δ score measures the magnitude and direction of motif perturbation caused by the variant. Motif gain and loss events were defined by comparing motif bindings between the two allele-specific sequences. A motif gain event was defined when a motif binding site was significant in the alternative allele sequence but not significant in the observed allele sequence:

$$\text{ScoreUbs} \geq \text{Tbind and ScoreObs} < \text{Tbind}.$$

On the other hand, a motif loss event was defined when a motif binding site was significant in the observed allele sequence but not significant in the alternative allele sequence:

$$\text{ScoreObs} \geq \text{Tbind and ScoreUbs} < \text{Tbind}.$$

Because multiple motif hits may occur within the sequence window for a given motif, we reduced the results to at most one gain event and one loss event per (variant, motif) pair. Specifically, the gain event with the maximum ΔMotif was retained, and the loss event with the minimum ΔMotif was retained. Ranked by ΔRegulatory Activity predicted by the BlueSTARR model, the top 100,000 variants (top) and bottom 100,000 variants (low) were chosen. The top variant set represents variants that strongly shift the predicted regulatory activity. The low variant set represents background variants as their ΔRegulatory Activity are merely zero. To identify transcription factor motifs preferentially perturbed among prioritized variants, motif gain and loss frequencies were compared between the top and low variant sets.

To control for sequence composition bias, the same motif perturbation analysis was also performed on dinucleotide-shuffled sequences. The dinucleotide-shuffled control was generated for both top and low variant sets. This produced four conditions: Top (Observed), Top (Shuffled), Low (Observed), and Low (Shuffled). For each motif, we counted the number of variants exhibiting at least one motif gain or loss event under each condition. Binary indicators were defined for each variant-motif pair:

- $G_i$ denotes the variant group indicator (Top vs Low)
- $S_i$ denotes the sequence type indicator (Observed vs Shuffled)

The probability that variant i exhibits a motif perturbation for motif m was modeled as:

$$\text{logit}(p_{im}) = \beta_{0m} + \beta_{1m} G_i + \beta_{2m} S_i + \beta_{3m} (G_i S_i)$$

The interaction term $\beta_{3m}$ tests whether motif perturbation enrichment in the Top variant set is stronger in observed sequences than expected from sequence composition alone. Exponentiating this coefficient yields the ratio of odds ratios comparing observed sequences with dinucleotide-shuffled controls.

$$\exp(\beta_{3m}) = \frac{\mathrm{OR}^{\mathrm{Observed}}_{\mathrm{Top:Low}}}{\mathrm{OR}^{\mathrm{Shuffled}}_{\mathrm{Top:Low}}}$$

For each motif, the null hypothesis H0: $\beta_{3m} = 0$ was tested using a Wald test derived from the logistic regression model. Gain and loss events were modeled separately. p-values were corrected for multiple testing using the Benjamini–Hochberg procedure to control the false discovery rate across motifs.

### S2. Supplementary Figures

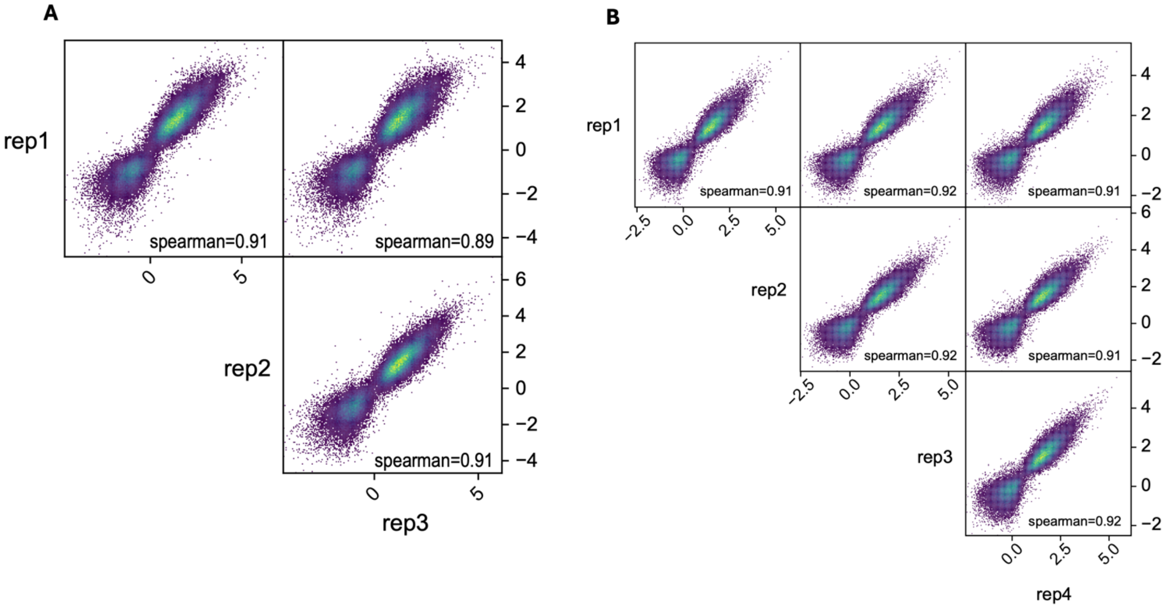


**Figure S1** Between replicate correlations for the input STARR-seq data. (A) K562 STARR-seq data between replicate correlations for the 3 DNA replicates. (B) A549 STARR-seq data between replicate correlations for the 5 DNA replicates.

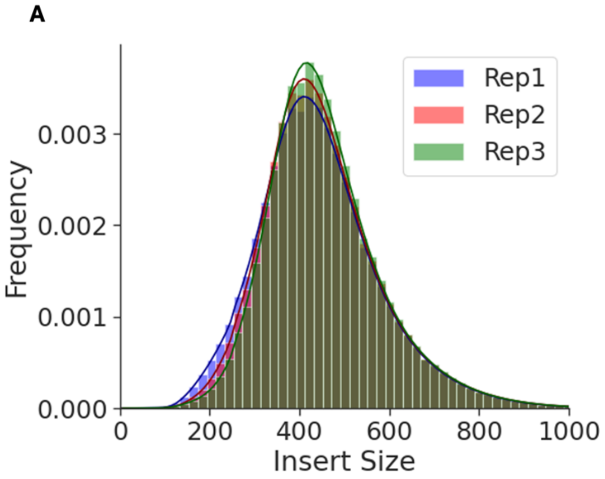


**Figure S2** Distribution of Insert sizes in the input data. (A) Distribution of insert sizes in the input library for K562 STARR-seq data. The mean insert size across the three DNA replicates was 451 bp. (B) Distribution of insert sizes in the input library for A549 STARR-seq data. The mean insert size across the 5 DNA replicates was 961 bp.

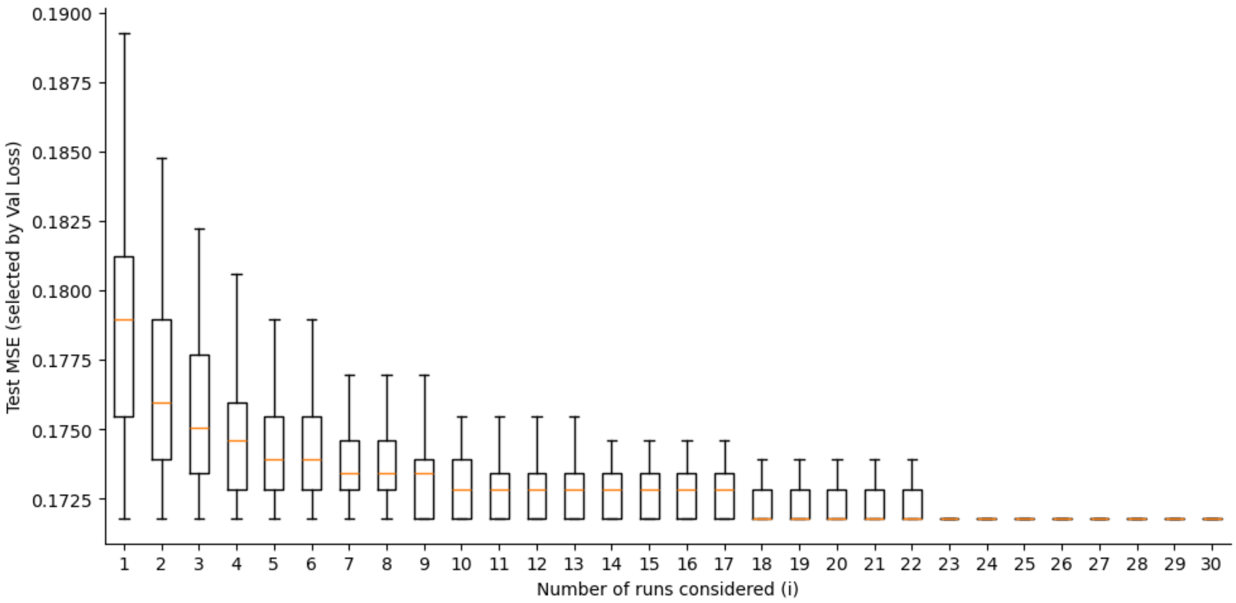


**Figure S3**. Distribution of Test MSE vs. Number of Runs under andomized run order. Each training run is independently initialized, so the best performing model may occur at any run index. To remove order effects, runs are randomly permuted 100 times. For each model i, the model with the lowest validation loss among the first i runs are selected and its test MSE is recorded. Boxplots show the resulting distribution, demonstrating convergence toward the optimal model as the number of runs increases.

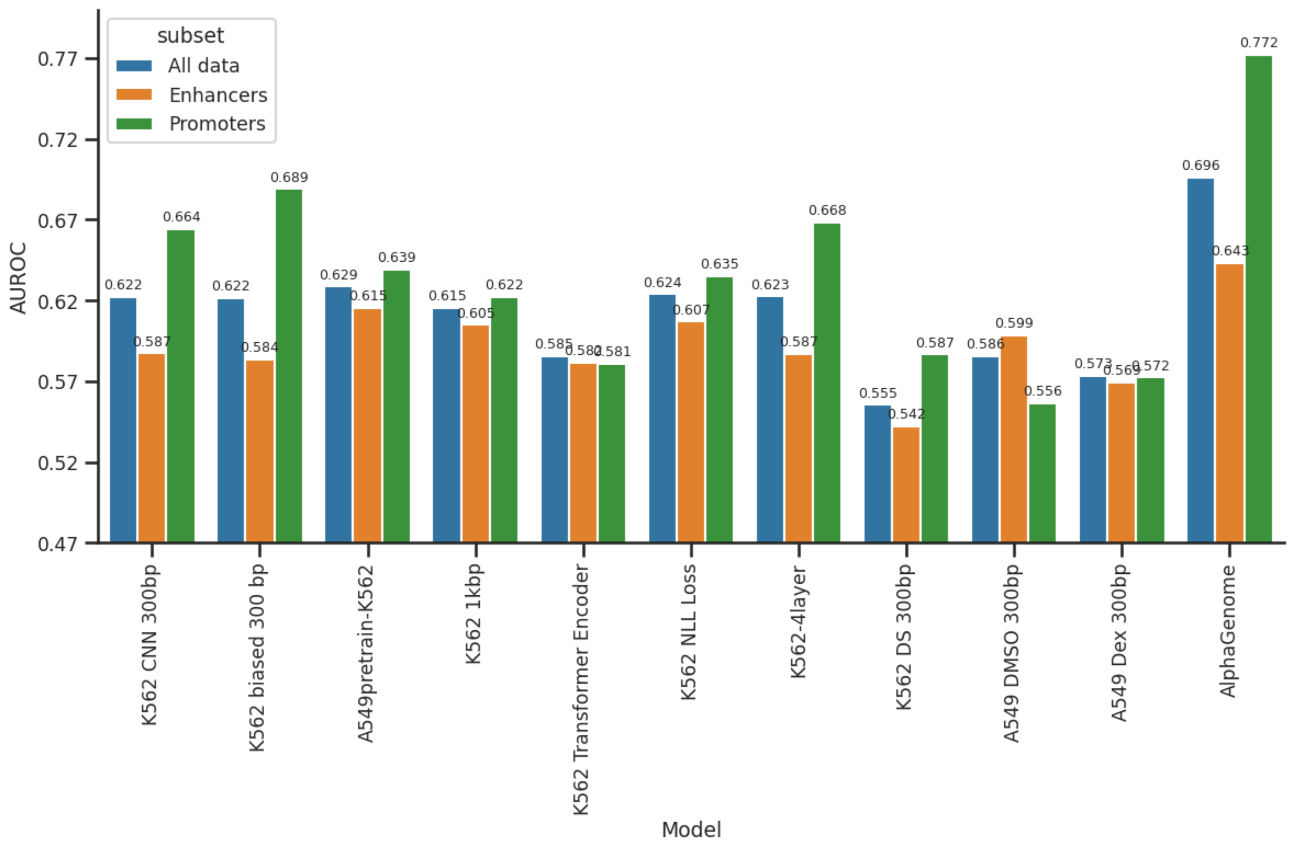


**Figure S4** AUROC on Kircher et al. data. The AUROC barplots for different versions of the BlueSTARR are benchmarked against Kircher et al., [4] MPRA saturation mutagenesis data for ~30000 variants. AUROC calculations were performed and visualized on all data and then separated into enhancers and promoters.

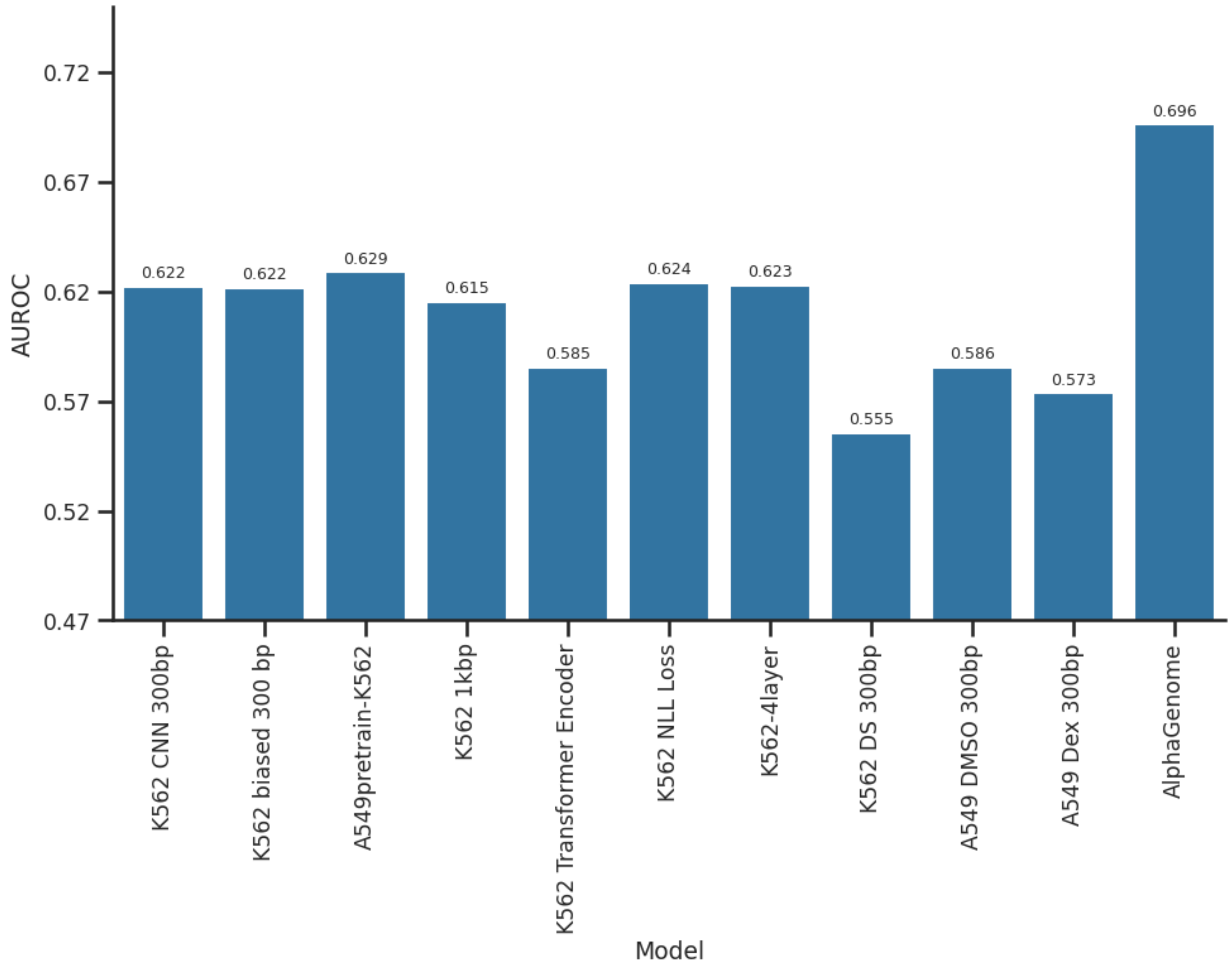


**Figure S5** AUROC on Kircher et al. data. The AUROC barplots for different versions of the BlueSTARR are benchmarked against Kircher et al.,[4] MPRA saturation mutagenesis data for ~30000 variants.

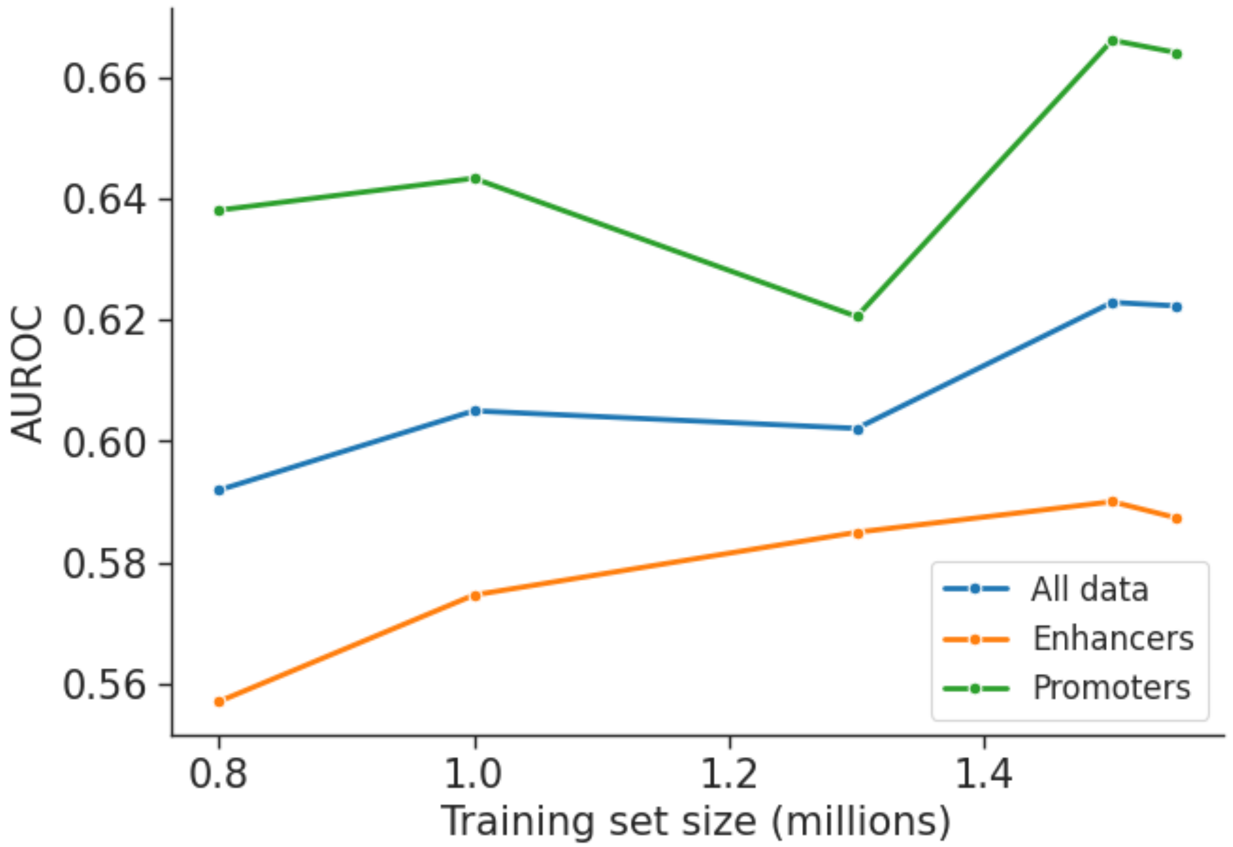


**Figure S6** Variation of AUROC with number of training examples. AUROC for K562 BlueSTARR MSE based loss model benchmarked against the Kircher et al. [4] saturation mutagenesis MPRA data for training set sizes varying from 0.8M to 1.55M. The plot also represents the variation of AUROC for training set size for enhancer and promoter elements along with all data.

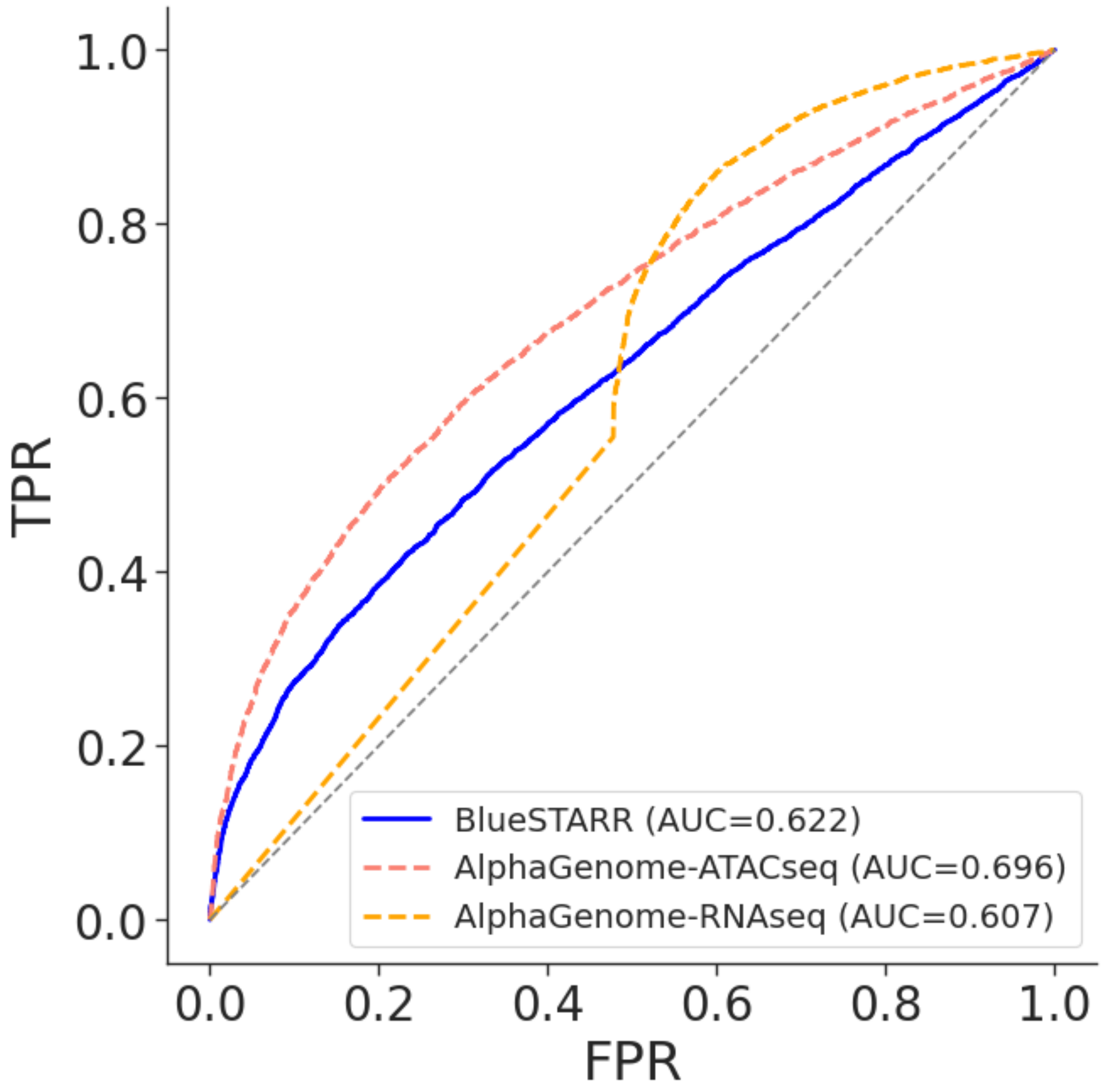


**Figure S7.** Comparison of AUROC between BlueSTARR (blue: AUC=0.622), Alphagenome ATAC-seq modality (orange: AUC=0.696)and Alphagenome RNA-seq modality (yellow: AUC=0.607) benchmarked against the Kircher et al. saturation mutagenesis MPRA data. AlphaGenome's training set included the test regions, possibly leading to data leakage. Missing values in AlphaGenome-RNAseq were replaced with zeros, leading to a non-smooth ROC.

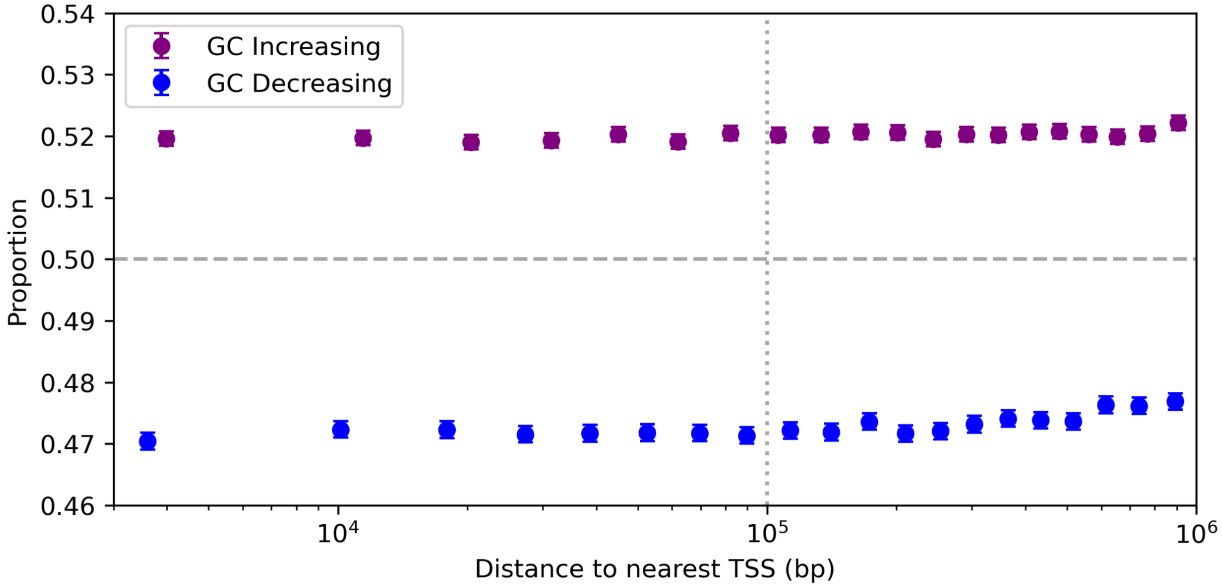


**Figure S8.** Distance-dependent enrichment of observed alleles in closed regions stratified by GC-content change. Figure 7 shows the distance-dependent pattern when GC-balanced mutation classes are aggregated (i.e., observed allele pairs containing exactly one G or C). Here, these cases are further stratified by mutation direction. Purple points correspond to GC-increasing mutations (reference A/T to SNV C/G), while blue points correspond to GC-decreasing mutations (reference G/C to SNV A/T). Points represent the proportion of sites at which an observed allele occupies the maximum predicted regulatory configuration within each distance bin, with 95% confidence intervals. GC-increasing and GC-decreasing mutation classes exhibit opposite baseline enrichment patterns but similar distance-dependent trends, indicating that the aggregate behavior reflects a mixture of directional mutation effects rather than a single homogeneous process.

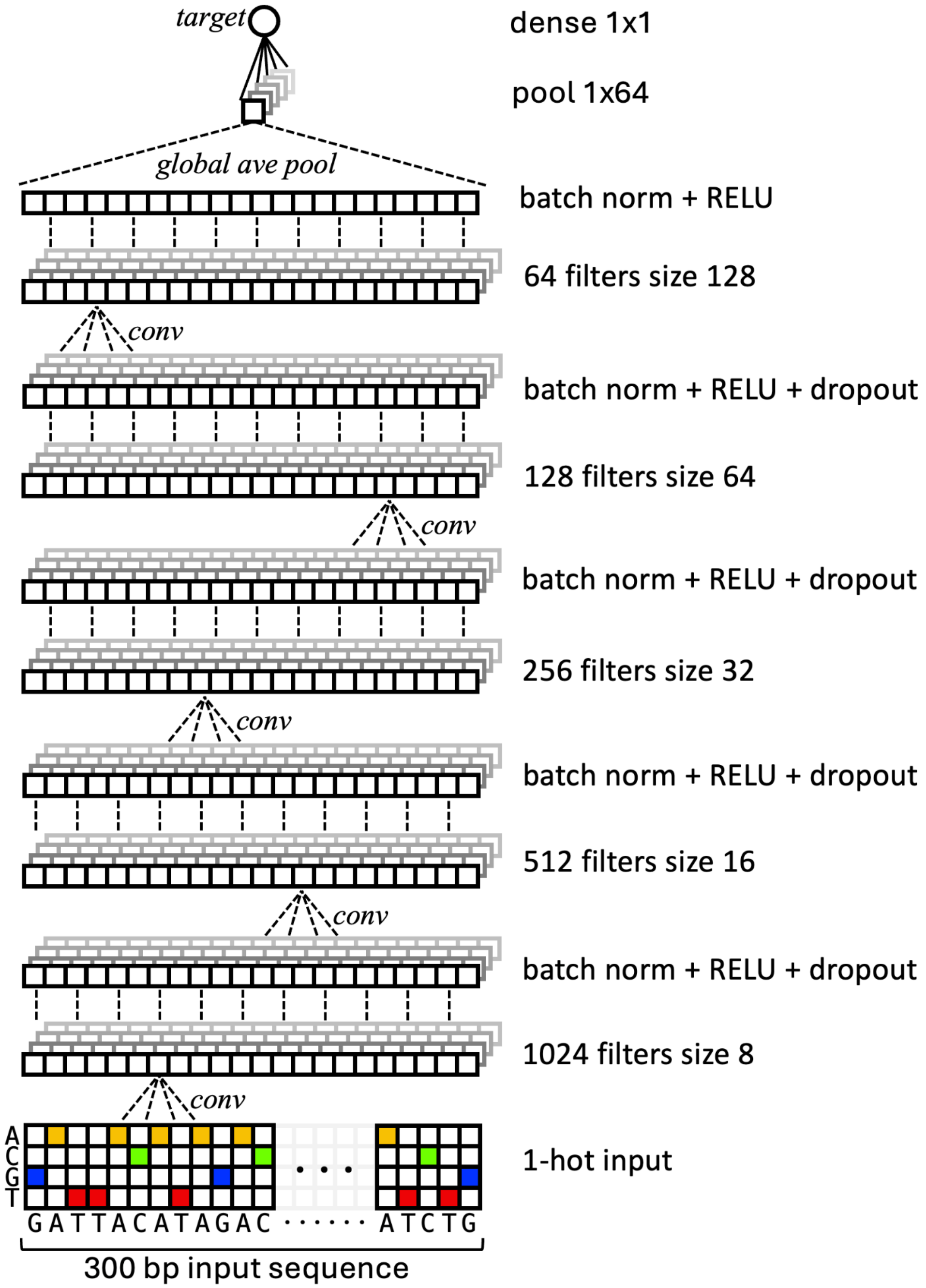


**Figure S9.** Default architecture of BlueSTARR used for most analyses in the main text, consisting of convolutional layers, RELU activation, dropout, and normalization, with a final global average pooling. Common architecture changes such as numbers of layers, numbers of filters and sizes, and whether to include additional blocks such as transformer encoder blocks, dense layers, pooling, and dilation can be specified in an editable configuration text file.

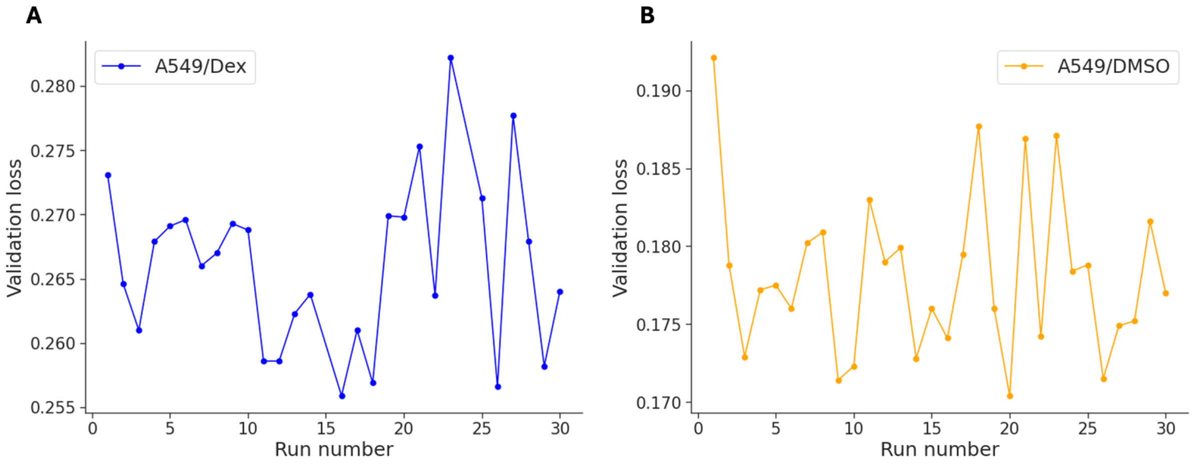


**Figure S10.** Model validation loss for each training run of the A549 DEX and A549 DMSO models. 30 independent training runs were initialized at the same time for both these models to account for stochasticity in model initialization and optimization.

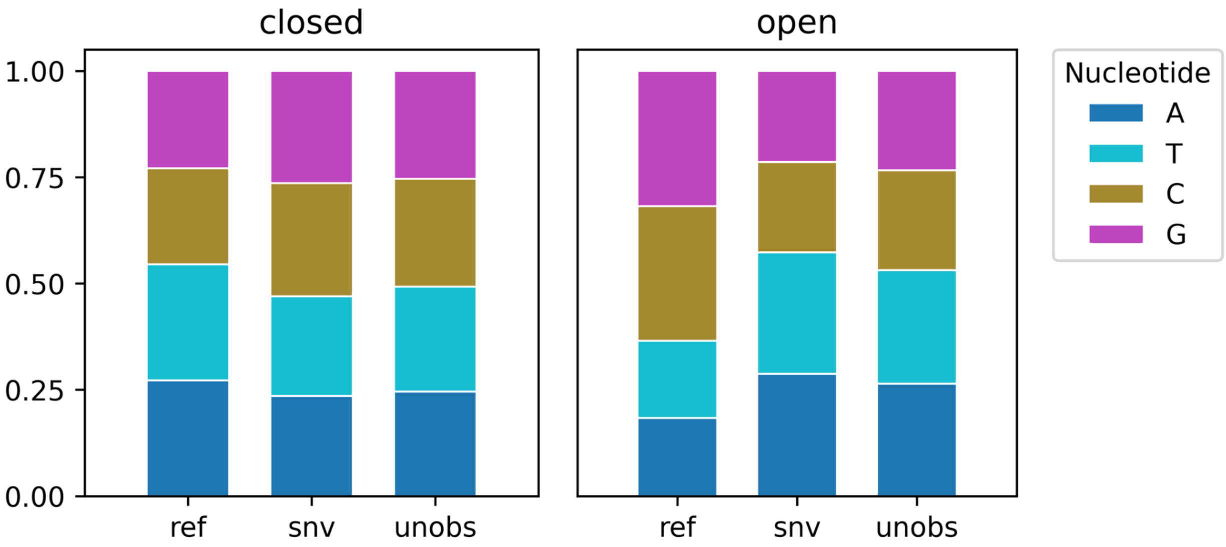


**Figure S11.** Stacked bar plots show the distribution of nucleotide identities (A, T, C, G) for reference (reference (`ref`), observed SNV (`snv`), and unobserved (`unobs`) allele classes in closed and open genomic regions. The composition of nucleotides differs across allele classes in both datasets, indicating that observed and unobserved allele groups are not compositionally matched. These differences motivate analyses that account for nucleotide-specific biases when comparing observed and unobserved alleles.

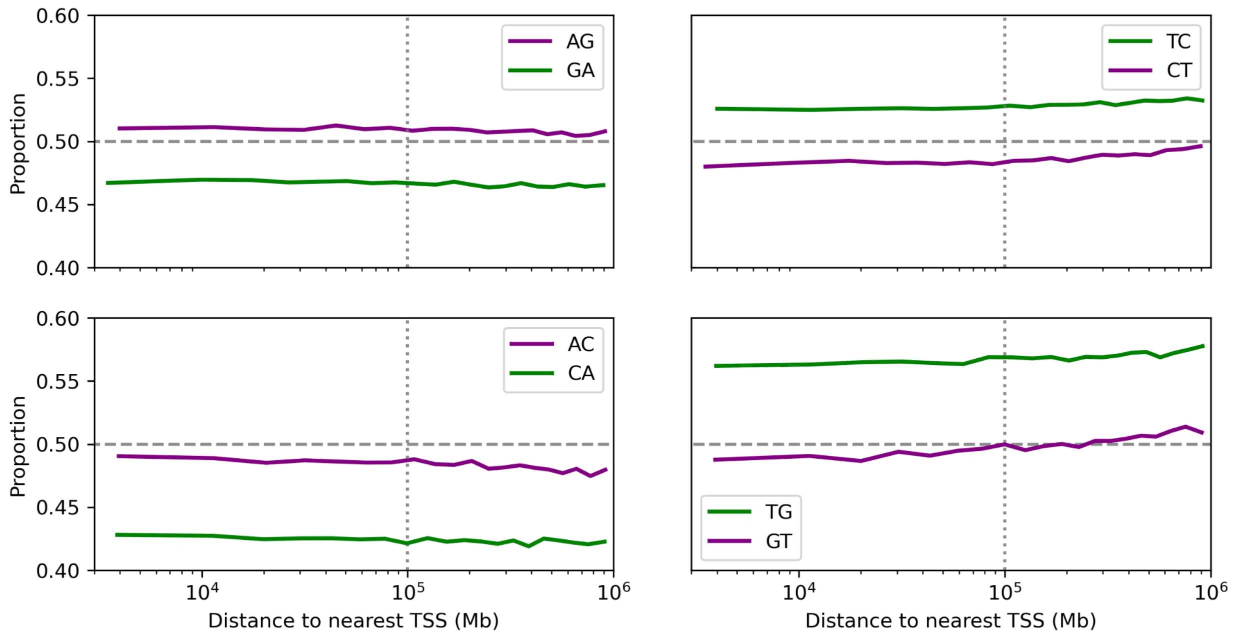


**Figure S12.** Distance-dependent enrichment patterns for GC-balanced mutation classes (i.e., observed allele pairs containing exactly one G or C) are shown after further stratification by mutation direction. Each panel displays the proportion of sites at which an observed allele occupies the maximum predicted regulatory configuration, binned by TSS distance quantiles. Lines correspond to directional mutation pairs (e.g., AG vs. GA). While GC-balanced mutation classes exhibit similar aggregate trends (Figure S8), directional mutation pairs show distinct baseline enrichment levels, indicating that mutation direction contributes substantially to the observed heterogeneity in enrichment patterns.

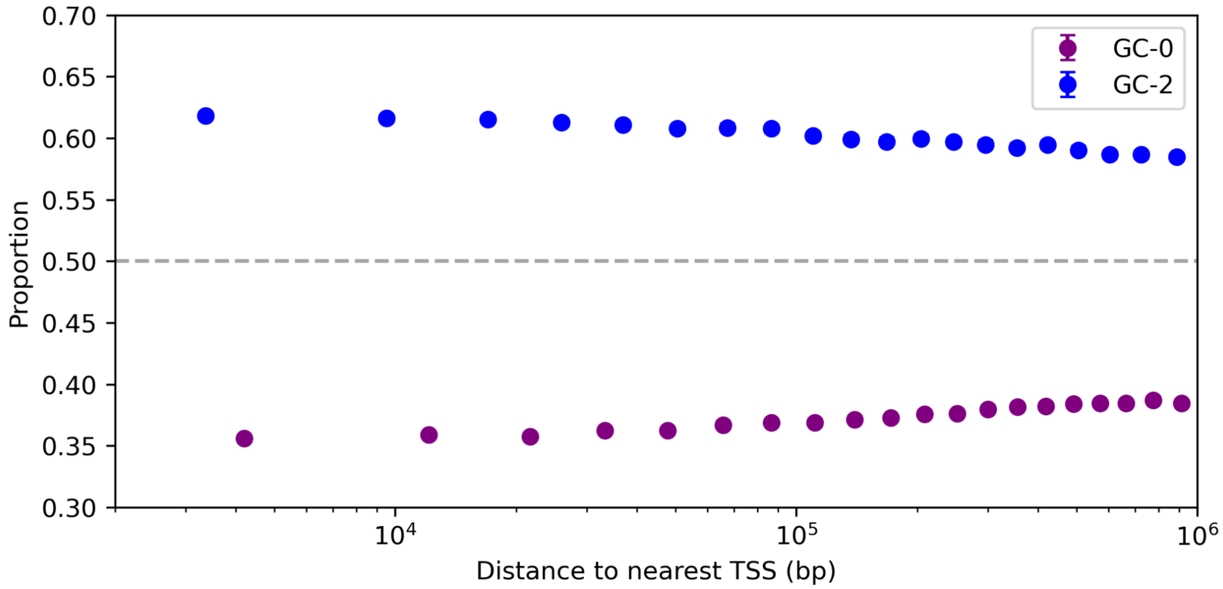


**Figure S13.** Distance-dependent enrichment for GC-0 and GC-2 mutation classes in closed regions. Proportion of sites at which an observed allele occupies the maximum predicted regulatory configuration is shown as a function of distance to the nearest transcription start site (TSS), binned by distance quantiles. GC-0 cases (purple) correspond to mutation pairs (A,T) and (T,A), while GC-2 cases (blue) correspond to (G,C) and (C,G). Points represent bin-level proportions. GC-0 and GC-2 mutation classes exhibit distinct baseline enrichment levels and modest distance-dependent trends, further demonstrating that enrichment patterns depend strongly on mutation class and nucleotide composition.

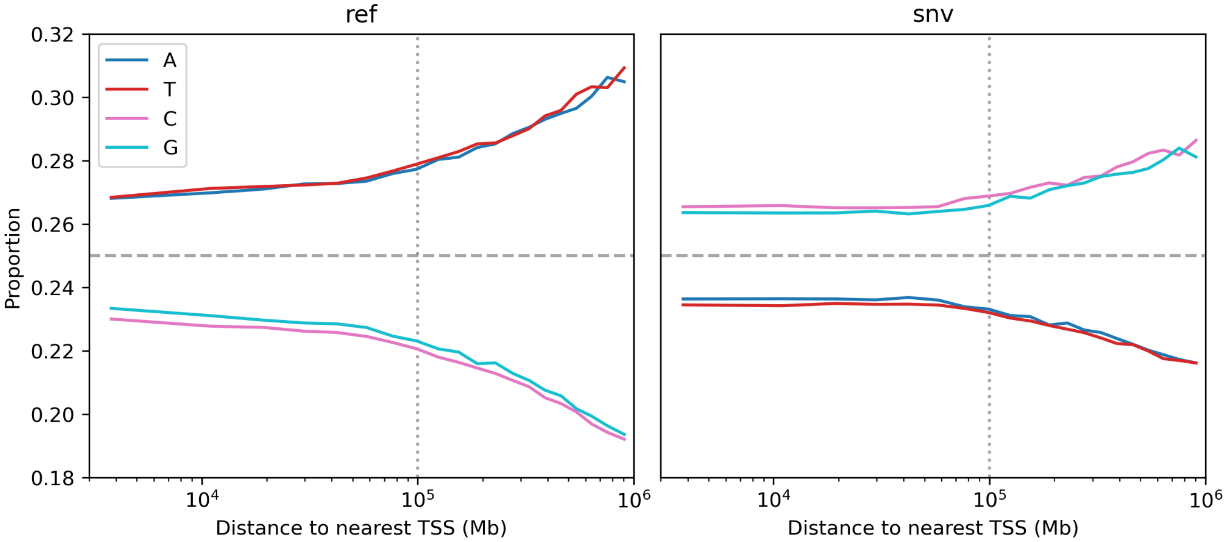


**Figure S14.** Nucleotide composition as a function of distance to the nearest TSS in closed regions. Proportions of nucleotide identities (A, T, C, G) are shown for reference alleles (left) and observed SNVs (right) across distance bins defined by TSS distance quantiles. The dashed horizontal line indicates the expectation under uniform nucleotide assignment (0.25). In reference alleles, A and T are overrepresented and increase modestly with distance, while C and G decrease. In contrast, SNVs exhibit the opposite pattern, with C and G increasing and A and T decreasing with distance. These opposing trends indicate that nucleotide composition varies with distance and differs systematically between reference and variant alleles, providing additional evidence that enrichment patterns may be influenced by nucleotide-dependent effects.
